# Integrated mesenchymal and extracellular cues drive bioengineered liver tissue formation and function

**DOI:** 10.1101/2025.08.28.672087

**Authors:** Shicheng Ye, Zhenguo Wang, Nalan Liv, Luc van der Laan, Jos Malda, Bart Spee, Frank G. Van Steenbeek, Kerstin Schneeberger-Verjaal

## Abstract

Human liver tissue engineering holds promise for creating physiological *in vitro* models but faces challenges replicating liver complexity. In the present study, we created bioengineered liver tissues (BLTs) utilizing three different cell types; human intrahepatic cholangiocyte organoids (ICOs), hepatic stellate cells (HSCs), and mesenchymal stromal cells (MSCs). Co-culturing with HSCs and MSCs accelerated growth and spontaneous fusion, resulting in complex liver-like tissue structures. In a dynamic suspension culture, BLTs had a more compact morphology and higher expression of hepatic markers, including *ALB*, *CYP3A4*, and *MRP2*. We further showed that animal-derived Matrigel can be replaced by a synthetic polyisocyanide (PIC)-based hydrogel for BLTs. Importantly, PIC-based hydrogel further promoted the maturation of BLTs assessed by parameters as intracellular protein levels, morphological analysis, and metabolic activity. Transcriptomic analyses revealed mechanisms underlying tissue formation and function. To conclude, our strategy yields functional liver tissues suitable for disease modelling, drug screening, and toxicity tests, and forms an important basis for future development of larger liver tissues for *in vivo* transplantation.

## Introduction

Organoids have emerged as powerful tools for disease modeling and fundamental biological studies, offering a promising platform to replicate key aspects of human tissue architecture and function. However, epithelial liver organoids lack the cellular and structural complexity necessary to accurately mimic liver physiology. This limitation hampers their utility in dissecting the roles of individual cell types and extracellular matrix (ECM) components in liver homeostasis, disease progression, and regeneration. Combining organoids with non-parenchymal cell types and defined matrices into bioengineered liver tissues (BLTs) will allow for models that more faithfully recapitulate native liver function and provide insights into the mechanistic processes governing tissue organization and maturation.

The fundamental components of BLTs are functional hepatic cells and supportive biomaterials that function as extracellular matrix (ECM) mimicry. The human liver consists of approximately 70-80% parenchymal cells (mostly hepatocytes and 3-5% cholangiocytes) and 20-30% nonparenchymal cells, mainly including 5-10% mesenchymal cells, 10-15% endothelial cells, around 3% Kupffer cells, and lymphocytes (*1–3*). Hepatocytes are the major cell type that performs vital liver functions; however, due to the limited availability of primary human hepatocytes (PHH), current tissue engineering strategies have shifted to stem cell-based cell sources and their subsequent differentiation into hepatocyte-like cells (*4*, *5*). Human intrahepatic cholangiocyte organoids (ICOs) mainly consist of adult liver stem cells, can be expanded for a long time into a large number of cells with stable genomic profiles, and can also be differentiated into hepatocyte- and cholangiocyte-like cells *in vitro*, making them suitable for LTE (*6–8*). However, since ICOs consist of epithelial cells only, they are far from mimicking the complexity and function of the liver. For example, mesenchymal cells, including liver mesenchymal stromal cells (MSCs) and hepatic stellate cells (HSCs), play important roles in liver regeneration and directly impact liver function (*9*, *10*). They secrete growth factors, such as hepatocyte growth factor (HGF), epidermal growth factor (EGF), fibroblast growth factor (FGF), transforming growth factor-beta (TGFβ), and bone morphogenetic protein (BMP), produce and remodel the ECM, and regulate the stem cell niche (*1*, *11*). All these factors influence parenchymal cells in liver homeostasis, as well as in liver regeneration. Importantly, mesenchymal cells also enable tissue formation *in vitro* and improve engraftment after transplantation (*12*, *13*). Here, we evaluated the use of bi-potential epithelial organoids (ICOs, hereafter organoids) and mesenchymal cells (HSCs and MSCs) for the generation of BLTs.

Various hydrogels have been developed to mimic the ECM for TERM approaches, both *in vitro* and *in vivo* (*14*, *15*). For organoid culture, Matrigel is currently the gold standard, primarily due to its complex biological composition, which supports organoid formation and proliferation. Due to shortcomings of Matrigel, such as being animal-derived, ill-defined, and with large batch-to-batch variations, scientists have developed (semi-)synthetic hydrogels as Matrigel alternatives, although the biological complexity of Matrigel is hard to mimic (*14*, *15*). Among those (semi-)synthetic hydrogels, polyisocyanide-based (PIC) hydrogels are drawing increasing attention in the field of TERM (*16*, *17*). PIC is synthetic and thermoreversible and was originally developed to mimic the porous and fibrous architecture and mechanical properties of ECM proteins, such as collagen and fibrin (*18–20*). With these advantages, PIC hydrogels have been successfully used for organoid culture *in vitro* (*16*, *17*) and for *in vivo* wound healing studies (*21*, *22*).

In this study, we bioengineered liver tissues by co-culturing human liver epithelial organoids with mesenchymal cells, including HSCs and MSCs. We cultured BLTs on an orbital shaker to create a dynamic microenvironment. Dynamic suspension (DS) culture methods have been previously reported to accelerate the growth of human epithelial organoids (*8*, *23*) and/or promote the maturation of organoids (*24–26*). In our co-cultures, the DS method enhanced spontaneous tissue formation. Subsequently, we replaced Matrigel with a well-defined hydrogel for the co-culture, using the synthetic, thermoreversible PIC supplemented with type 1 collagen. We investigated tissue formation, cellular composition, morphology, and maturation of the BLTs in all conditions. Further, we conducted transcriptomic analyses of BLTs and propose a model explaining the contributions of cellular and extracellular factors to tissue formation and maturation. As such, this study provides a straightforward method for bioengineering complex human liver tissues that can be directly applied for studying cell-cell and cell-matrix interactions in homeostasis, disease and regeneration.

## Results

### Mesenchymal Cells Enhance the Expansion of Organoids

The human liver consists of multiple types of cells, and missing any cell type leads to liver dysfunction. To mimic this cellular complexity in bioengineered human liver tissues (BLTs), we co-cultured organoids with HSCs and MSCs in expansion medium (EM) at a 6:1:1 ratio (**Figure 1A**). We observed that all co-cultures formed more and larger organoids than the control group with organoids only 4-7 days (D4-D7) after single-cell seeding in Matrigel (**Figure 1B**). We confirmed this observation by quantification of organoid numbers (**Figure 1C**) and organoid diameters (**Figure 1D**). On day 11 (D11), the co-cultures were almost confluent while organoids in the control group were sparser (Figure 1B). When we characterized the co-cultures with qPCR assays, we found that the stem cell/progenitor marker *LGR5* was significantly lower in co-cultures containing HSCs (**Figure 1E**). Hepatocyte markers, including *HNF4A*, *ALB*, and *CYP3A4,* indicated that all cultures expressed the markers at similar levels, which were much lower compared to liver tissue, in line with expansion conditions. Importantly, the expression of the ductal marker *KRT19* was comparable among all groups, indicating that HSCs and MSCs did not hamper the ductal phenotype of organoids cultured in EM. Instead, the mesenchymal cells enhanced the growth of organoids in EM. Controls containing only mesenchymal cells (both HSCs and MSCs) showed no obvious proliferation till D11 in EM (**Figure S1A**) and very low expression of hepatocyte marker genes, except for *BSEP* (**Figure S1B**).

**Figure 1.**
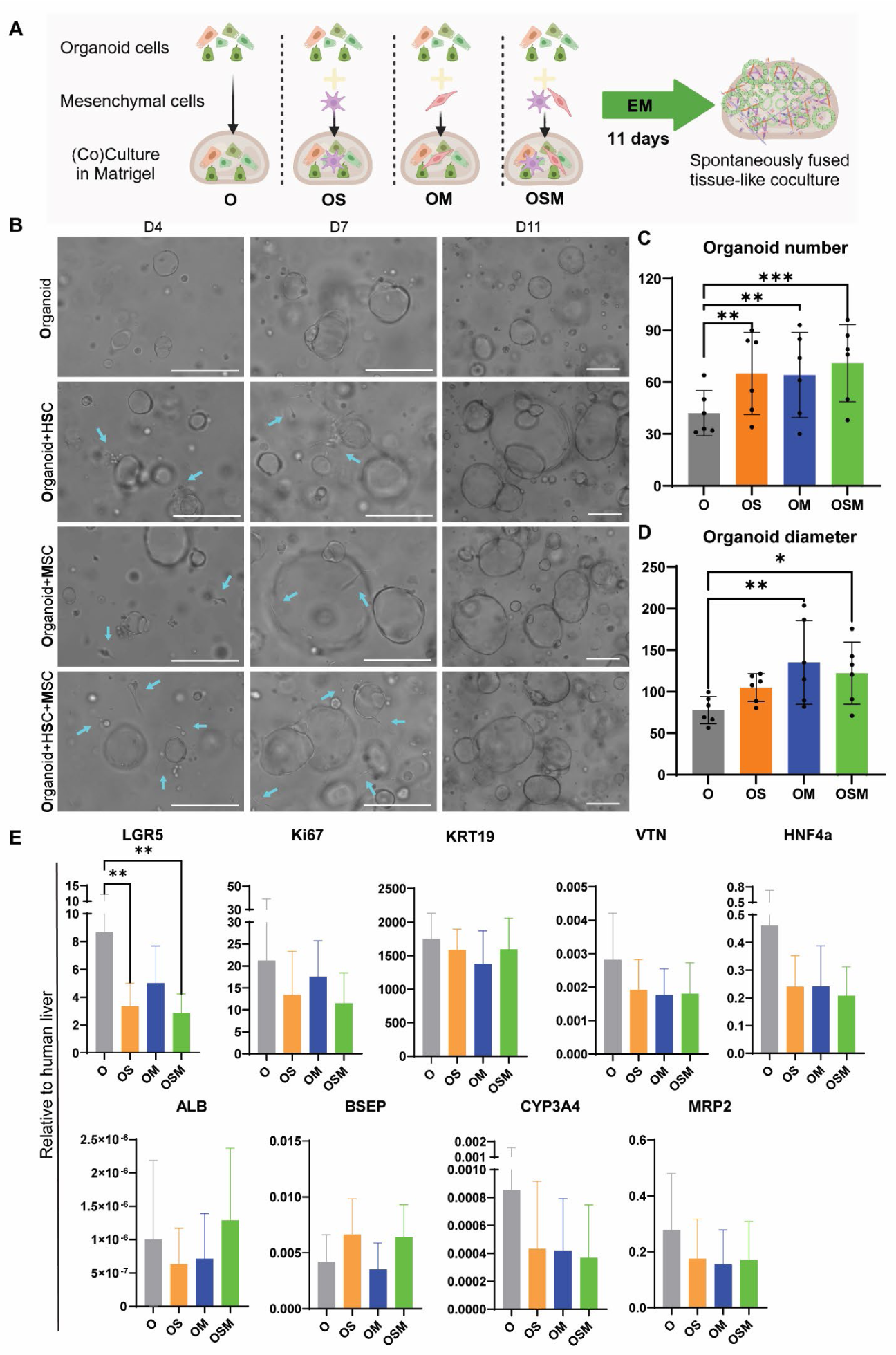
Expansion of co-cultures in Matrigel. n=6. (A) Schematic of different cellular compositions in Matrigel droplets. O for organoid, S for hepatic stellate cell (HSC), and M for mesenchymal stem cell (MSC). (B) Bright-field pictures showing the morphology of organoids co-cultured with or without HSCs and MSCs in Matrigel in expansion medium (EM) at three time points, D4 (left), D7 (middle), and D11 (right). Organoids only (O), organoids with HSCs (OS), organoids with MSCs (OM), organoids with HSCs and MSCs (OSM). Arrows indicate mesenchymal cells (MSC or HSC). Scale bar = 200 µm (C) Organoid number quantification. Organoid numbers were quantified from 6 donors and at least 3 pictures per donor. (D) Organoid diameter quantification. Organoid diameters were measured from 6 donors and at least 10 organoids per donor. (E) Gene expression of co-cultures expanded in Matrigel for 11 days, compared to human liver tissues. Stem cell/progenitor marker LGR5, proliferative marker Ki67, ductal marker KRT19, and hepatocyte markers HNF4a, ALB, CYP3A4, BSEP, MRP2, and VTN were used for the qPCR assays. Significant analysis was labeled with * (p<0.05).

### HSCs, but not MSCs, Hamper the Differentiation of Organoids

After establishing the co-cultures of organoids with HSCs and/or MSCs in expansion medium, we continued to induce hepatic differentiation with differentiation medium (DM) when the cultures were confluent (**Figure 2A**). After 8 days of differentiation, light microscopy analysis showed that organoids became more condensed and darker compared to EM and all the co-cultures fused into more complex structures (**Figure 2B**). Most HSCs and MSCs were not visible anymore as single cells like in EM conditions (Figure 1B), indicating that they may have fused with organoids. Organoids that had fused with surrounding cells seemed to have opened their lumen; however, this spontaneous fusion was not complete due to partial attachment of cells to the standard culture plates. In control groups containing only mesenchymal cells, we observed enhanced proliferation compared to EM conditions, with clear cellular interactions and MSC cluster formation (**Figure S2A**). Compared to the expression levels of EM conditions (Figure 1C), most hepatocyte markers (*ALB*, *CYP3A4*, and *BSEP*) were upregulated in all conditions after differentiation, while the stem cell marker *LGR5* and the proliferation marker *Ki67* were downregulated (**Figure 2C**). Hepatocyte markers were comparably expressed in organoid-only culture and the co-culture of organoids with MSCs, but the expression of most markers in co-cultures containing HSCs was significantly lower (**Figure 2C**). In contrast, the expression levels of *Ki67* were significantly higher in co-cultures containing HSCs. Once mesenchymal cell-only controls were included in the graphs, we found that the expression levels of LGR5 and Ki67 in mesenchymal cells were upregulated in DM compared to their expression in EM, particularly in HSCs, while hepatocyte markers remained low in mesenchymal cells (**Figure S2B**). We further compared the different co-cultures with histological analysis by H&E staining. H&E results showed that all cocultured conditions displayed more complex structures than the organoid-only condition, with the tri-culture (OSM) being the most compact (**Figure 2D**). Since the co-culture condition of organoids with HSCs and MSCs resulted in efficient tissue formation and most closely reflects the cellular complexity of the liver, we continued with this OSM condition for our further studies.

**Figure 2.**
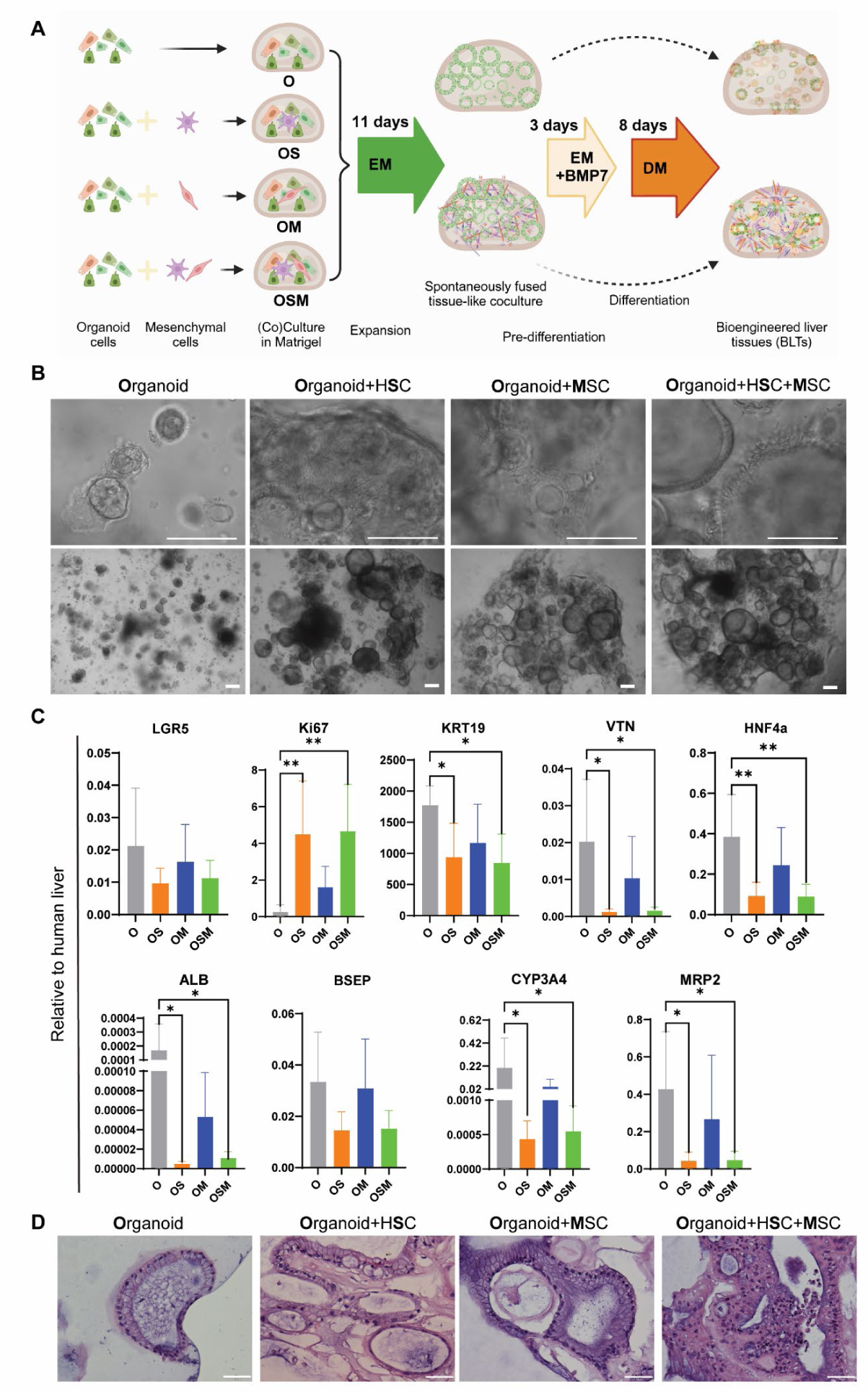
Differentiation of co-cultures in Matrigel under static culture (SC) conditions. n=6. (A) Schematic of different cellular compositions as co-cultures from EM to differentiation medium (DM). (B) Bright-field pictures showing the morphology of organoids co-cultured with HSC and/or MSCs in Matrigel with DM for 8 days at two magnifications, zoomed-in (top) and overall view (bottom). Scale bar=200 µm (C) Gene expression of co-cultures differentiated in Matrigel for 8 days, compared to human liver tissues. Primers of the same makers as in EM were used for the qPCR assays. Significant analysis is labeled with * (p<0.05). (D) H&E staining images showing the morphology of organoids co-cultured with HSC and/or MSCs in Matrigel with DM for 8 days. Scale bar=50 µm

### Dynamic Suspension Culture Promotes the Fusion and Differentiation of BLTs

To avoid attachment of cells to the bottom of the culture plates and enhance the spontaneous tissue formation, we next utilized low-attachment plates and transferred the plates to an orbital shaker for dynamic suspension culture (DS) during the differentiation process (**Figure 3A**). Bright-field pictures showed that the OSM-derived BLTs fused more than organoid-only derived BLTs in both static culture (SC) and DS. In addition, the OSM condition formed more compact tissues in DS conditions compared to SC (**Figure 3B**). Then, we compared the differentiation of all cultures by qPCR. Most hepatocyte markers (*ALB*, *CYP3A4*, *BSEP*, *MRP2*, and *MDR1)* showed a (trend of) higher expression in DS than in SC in organoid-only as well as OSM conditions (**Figure 3C**). Consistent with the previous results when differentiated in normal adhesion plates (Figure 2), BLTs derived from both OSM and organoid-only showed downregulation of stem cell marker *LGR5* and proliferation marker *Ki67* after differentiation compared to EM conditions (**Figure 3D**). Also, *Ki67* was significantly higher in BLTs derived from OSM than from organoid-only, along with a lower expression of the majority of hepatocyte markers (*CYP3A4*, *BSEP*, *KRT19*, *ECAD*, *HNF4a*, and *MDR1*) (**Figure 3C-D**). Interestingly, the extracellular matrix (ECM) component genes fibronectin (*FN*) and vitronectin (*VTN*) showed opposite trends of expression between organoid-only and OSM-derived BLTs, with *FN* higher in OSM and *VTN* higher in organoid-only cultures (**Figure 3D**). Moreover, the mechanotransduction marker *YAP* was significantly lower expressed in OSM than in organoid-only cultures, and the expression levels of *YAP* were comparable between SC and DS (**Figure 3D**), indicating differential expression of *YAP* in different cell types rather than differences in culture conditions. We further verified the hepatic and epithelial phenotype of both OSM and organoid-only derived BLTs on the protein level with immunofluorescent (IF) staining. IF results showed that all BLTs expressed Albumin (ALB) after differentiation, while no specific signal for CYP3A4 was observed (**Figure 3E**). The epithelial protein E-cadherin (ECAD) and the ductal protein KRT19 (K19) were both expressed in all conditions (**Figure 3F**).

**Figure 3.**
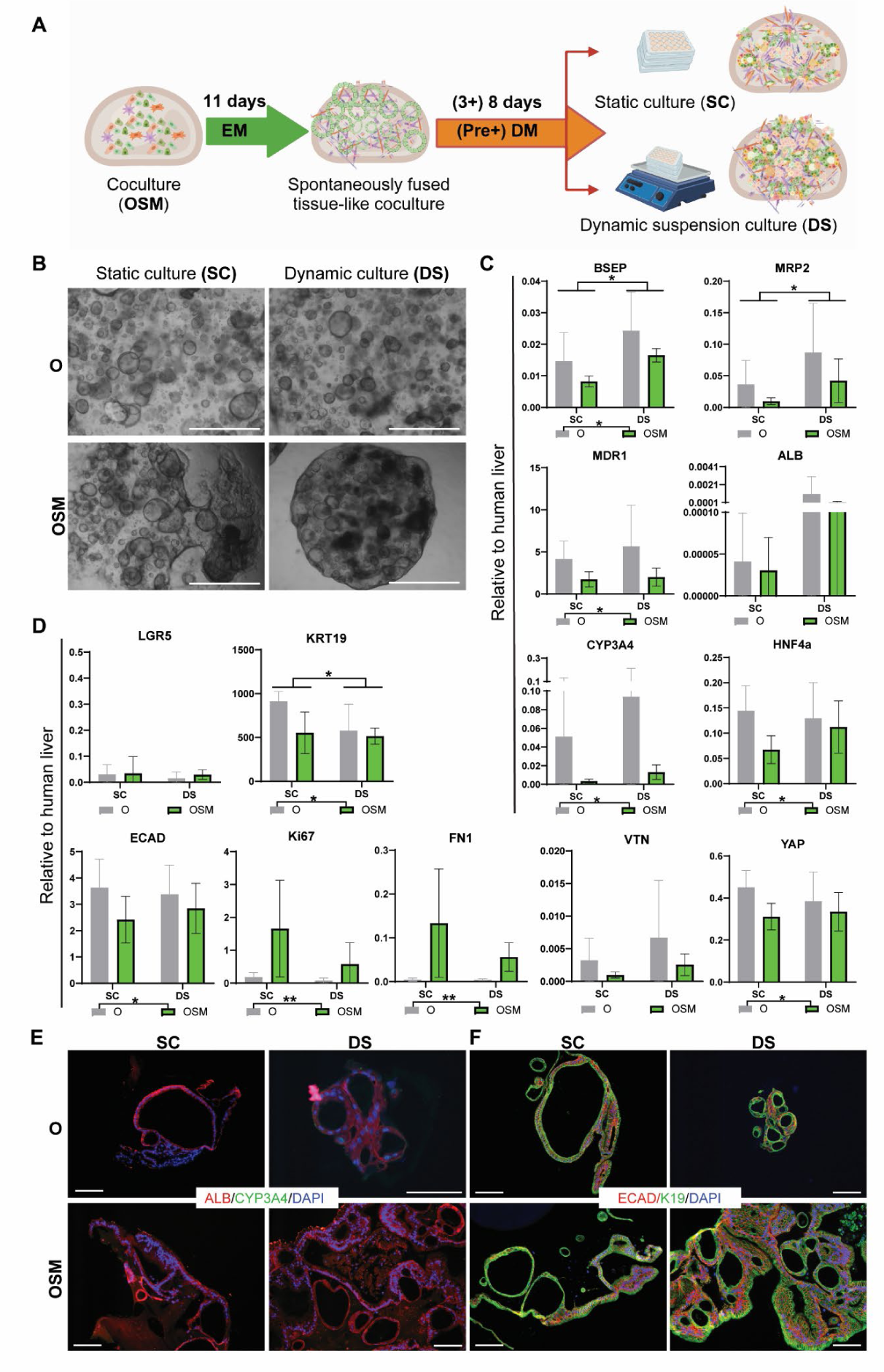
Characterization of BLTs in static or dynamic conditions. N=6. (A) Schematic of the experimental setup comparing static culture (SC) and dynamic suspension culture (DS) conditions. (B) Morphology of organoid-only (O) and co-cultures (OSM) derived bioengineered liver tissues (BLTs) in SC or DS conditions after 8 days of differentiation. DS meant the cells were plated in Matrigel droplets in low-attachment plates which were put on a horizontal shaker at a speed of 70 rpm. Scale bar = 1000 µm. (C-D) Gene expression of BLTs in SC vs DS conditions, compared to human liver tissues. Besides the markers previously used for SC characterization, more markers were used for the qPCR assays, including epithelial marker ECAD, polarity marker MDR1, ECM component FN, and mechanotransduction marker YAP. Significant analysis was labeled with * (p<0.05). (E-F) Characterization of BLTs by immunofluorescent (IF) staining. Functional hepatocyte proteins ALB (red) and CYP3A4 (green); Epithelial protein ECAD (red), and ductal protein KRT19 (K19, green). Scale bar =100 µm.

### PIC-Collagen Hydrogels Enhance the Maturation of BLTs

The co-culture of organoids with HSCs and MSCs formed fused liver tissues (BLTs) in Matrigel droplets with liver functionality after differentiation. However, Matrigel has many well-known disadvantages, such as large batch-to-batch variation and its mouse tumor origin, which make it unsuitable for clinical applications. In addition, we hypothesized that the immature nature of our BLTs was partly caused by the abundant and diverse ECM components in Matrigel that may maintain a stem cell niche and prevent further differentiation. Modulation of the microenvironment was shown to improve engraftment and function after transplantation (*5*). Hence, we developed polyisocyanide (PIC)-based hybrid-hydrogels to bioengineer liver tissue, supplemented with two major ECM proteins, either laminin-entactin complex (LEC) or collagen-1 (**Figure 4A**). Previously, we reported that PIC-LEC (PL) hydrogel was sufficient for the expansion of organoids, whereas organoid differentiation towards hepatocytes in PL was incomplete, comparable to Matrigel (*16*). Here, we supplemented PIC with collagen-1 to enhance hepatic differentiation (*27*). Consistent with the morphology in Matrigel droplets, OSM fused more than organoid-only culture and formed compact BLTs with the DS method, especially in the PC hydrogel (**Figure 4B**). Gene expression analysis by qPCR assays showed that most hepatocyte markers were significantly higher expressed in PIC hydrogels, particularly in PC, compared to Matrigel (data not shown). Then, we compared the differentiation of BLTs derived from organoid-only and OSM in PL and PC, respectively. The results showed that some of the hepatic markers analyzed (*ALB*, *BSEP*, and *MDR1*) were comparably expressed in PL and PC, whereas others (*CYP3A4*, *MRP2*, and *HNF4a*) were significantly upregulated in PC compared to PL (**Figure 4C**). Interestingly, the stem cell marker *LGR5* and proliferation marker *Ki67* were significantly higher expressed in PC than in PL, just as the ductal marker *KRT19*, ECM components *FN* and *VTN*, and mechanotransduction marker *YAP* (**Figure 4D**). The epithelial marker *ECAD* was well maintained in all conditions. We further verified the expression of hepatic markers on the protein level by IF staining. The results confirmed the presence of both ALB and CYP3A4 proteins (**Figure 4E**). Notably, CYP3A4 was not detected at protein level in BLTs differentiated in Matrigel droplets (Figure 3E). Epithelial protein ECAD and ductal protein KRT19 were both detected in all BLTs (**Figure 4F**). In summary, we showed that PIC hydrogels are suitable to replace Matrigel for liver tissue engineering using co-cultures (OSM) of organoids with HSCs and MSCs, and supplementation of PIC with collagen-1 improved fusion and hepatic maturation of the BLTs.

**Figure 4.**
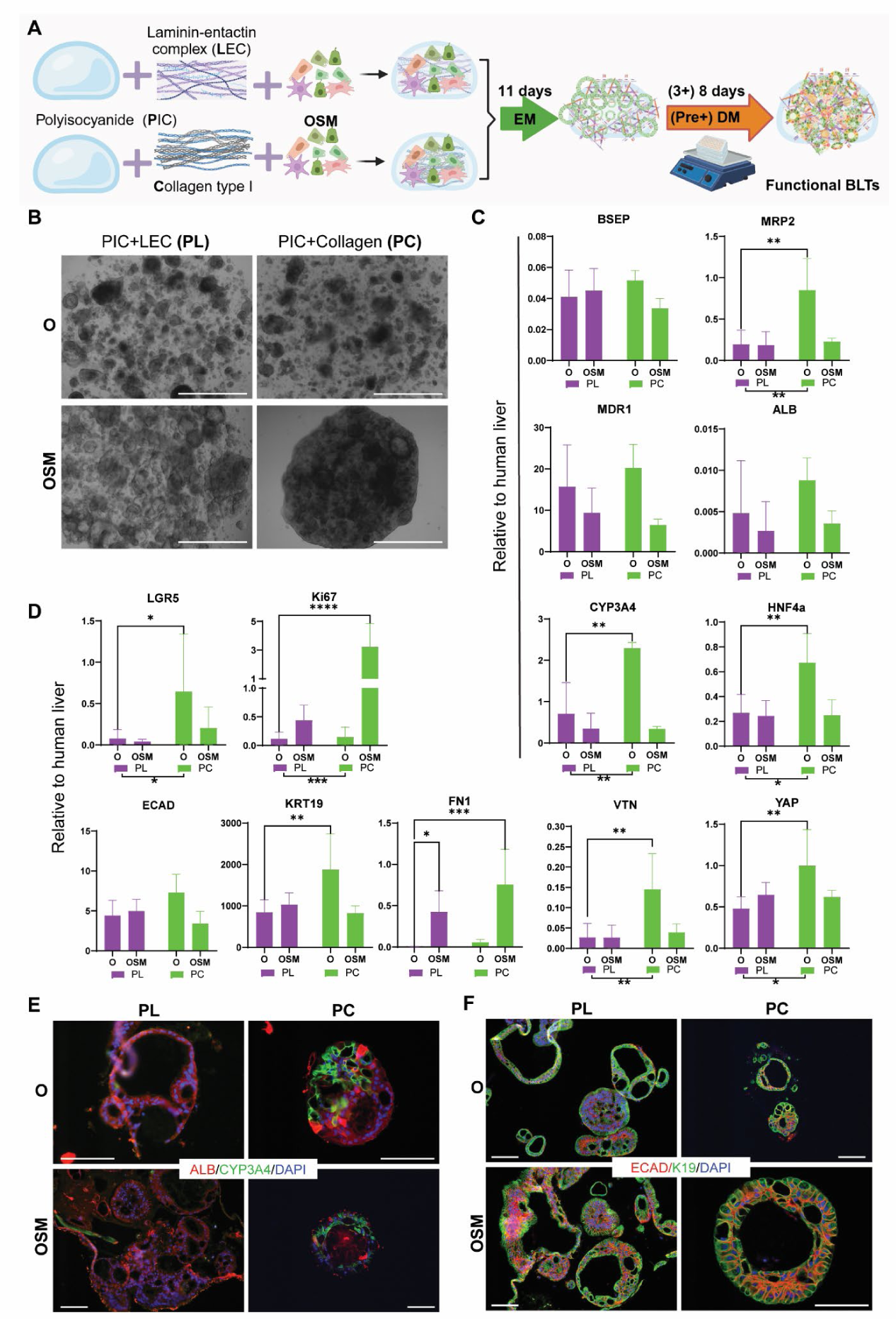
Characterization of BLTs formed in PIC hydrogels. n=3. (A) Schematic of co-cultures (OSM) in polyisocyanide (PIC)-based hydrogels for bioengineered liver tissues (BLTs). (B) Morphology of BLTs in PIC hydrogels after 8 days of differentiation. PL = PIC + Laminin-entactin complex (LEC); PC = PIC + Collagen-1. Scale bar =1,000 µm. (C-D) Gene expression of BLTs differentiated for 8 days in PIC hydrogels, compared to human liver tissues. The same markers were used as for characterizing samples in Matrigel under DS conditions. (E-F) Characterization of BLTs by IF staining. (E) Functional hepatocyte proteins ALB (red) and CYP3A4 (green); (F) Epithelial protein ECAD (red) and ductal protein KRT19 (green). Scale bar =100 µm.

### BLTs Retain Long-term Functional Hepatic Characteristics

We further investigated the hepatic functions of the BLTs with PC hydrogels after long-term maintenance (15 days) in dynamic suspension (DS). These include intracellular protein levels of ALB, ALAT, ASAT, GLDH, and ASAT in the culture medium (**Figure 5A-E**). We observed an overall decrease in ALB level from D8 to D15 for both organoid-only (O) and co-culture (OSM) conditions. Interestingly, there was no significant difference between O and OSM conditions anymore at D15 (Figure 5A). ALAT levels remained similar between D8 and D15, and again, we did not observe a significant difference between O and OSM conditions anymore at D15 (**Figure 5B**). The intracellular levels of ASAT between O and OSM were similar, as between D8 and D15 (**Figure 5C**). Interestingly, ASAT levels in the medium were always lower in OSM than O, with no difference between D8 and D15, indicating that there might be less cell damage in the OSM condition (**Figure 5D**). Contrary to the decreased ALB levels, there was a trend of increasing GLDH levels from D8 to D15, with no difference between O and OSM (**Figure 5E**). Consistent with intracellular GLDH levels, the ammonia elimination ability of BLTs increased from D8 to D15 with comparable levels between O and OSM conditions (**Figure 5F**). We then characterized the BLTs with histological analyses after 15 days of differentiation. H&E images show that there were overall more and bigger fused structures in OSM than in O (**Figure 5G**). Both O- and OSM-formed BLTs displayed remarkable glycogen storage, as determined by PAS staining (**Figure 5H**). Notably, organoids in Matrigel have a clear polarity with the apical side in the lumen and the basal side facing the matrix. However, based on the location of the nucleus and glycogen from the H&E and PAS stainings, we found that the polarity of organoids in the BLTs in PC could have changed. Therefore, we conducted IF staining with hepatocyte marker keratin 18 (K18), apical marker BSEP, and basolateral marker MRP3. The results show that the apical protein BSEP is located mostly on the outside of organoids in PC with clear presence of K18 and MRP3, which is strongly expressed in certain locations of the BLTs (**Figure 5I**). Further, we investigated the subcellular structures of BLTs with transmission electron microscopy (TEM) analysis. The TEM images show the presence of tight junctions, bile canaliculi-like structures, and microvilli in both O- and OSM-derived BLTs in PC (**Figure 5J**). Compared to BLTs in Matrigel (**Figure S3**), BLTs in PC formed more tight junctions and bile canaliculi-like structures, and their microvilli were located both outside of the organoids/BLTs and inside the lumens (**Figure 5J**), which is another sign of mixed polarity. Furthermore, we treated BLTs with a cocktail of substrates of phase I (CYP3A4, CYP1A2) and phase II (UGTs) enzymes and measured the parent compound depletion and metabolite formation with LC-MS/MS. The results show that both O- and OSM-derived BLTs in PC were able to perform comparable drug metabolizing capabilities after 15 days in culture (**Figure K-M**). Taken together, BLTs in PC formed structures closer mimicking natural liver than in Matrigel and retained key hepatic functions after long-term culture.

**Figure 5.**
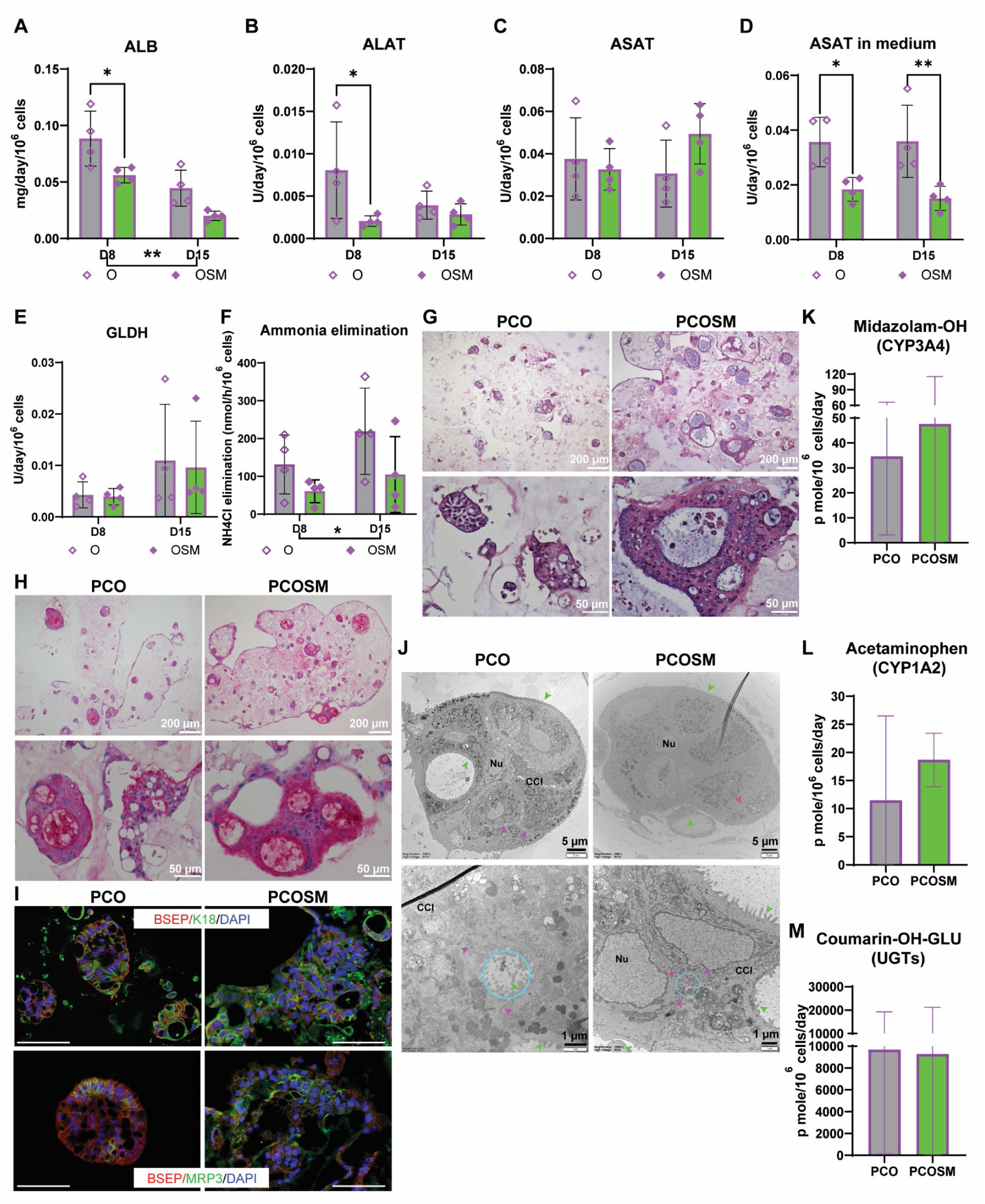
BLTs retain long-term functional hepatic characteristics. (A-C) Intracellular protein levels of ALB, ALAT, ASAT. (D) ASAT levels in the medium. (E) Intracellular levels of GLDH. (F) Ammonia elimination assay to assess the ammonia detoxification capacity of bioengineered liver tissues (BLTs). The assessment of protein levels was measured with medium incubated with cells for 24 hours. The BLTs for ammonia elimination assays were incubated with DM with the addition of NH4Cl for 24 hours. Both protein levels and ammonia elimination assays were assessed at two timepoints (i.e. D8 and D15). (G) H&E staining and (H) PAS staining of BLTs. (I) IF staining with apical marker BSEP in combination with hepatocyte marker keratin 18 (K18, upper) or with basolateral marker MRP3 (lower). Scale bar=100 µm. (J) Subcellular characterization of BLTs with TEM analysis. Nucleus (Nu), bile canaliculi-like structures (blue circle), tight junction (purple arrow), microvilli (green arrow), and cell-cell interactions (CCI). (K-M) Compounds assays to determine metabolizing capabilities of BLTs. Metabolite formation was measured by LC-MC/MS and normalized to live cell numbers, reflecting the enzymatic activities of phase I (CYP3A4, CYP1A2) and phase II (UGTs), displayed in (K), (L), (M), respectively. n=4

To better understand the molecular mechanisms driving tissue formation and maturation in the different culture conditions we used, we conducted bulk RNA-seq analyses. Principal component analysis (PCA) displays clear clustering of organoid-only (O) and co-culture (OSM) conditions (**Figure S4A**). Within OSM conditions, samples formed several small subgroups mainly based on hydrogels - Matrigel (MG) or PIC + Collagen (PC) - and donors (d1, d2, d3, d4). While, the differences between static culture (SC) and dynamic suspension (DS) conditions or between differentiation times (D8 vs D15) are even less clear than donor variations. Consistent with the PCA graph, Pearson correlation analysis shows similar clustering of all samples used for bulk RNA-seq (**Figure S4B**). To further investigate the major influences of different culture conditions, we conducted paired analysis within each condition, such as O vs OSM (**Figure 6, Figure S5**), MG vs PC (**Figure 7, Figure S6**), SC vs DS (**Figure S7**), and D8 vs D15 (**Figure S8-1** for O and **Figure S8-2** for OSM, respectively).

**Figure 6.**
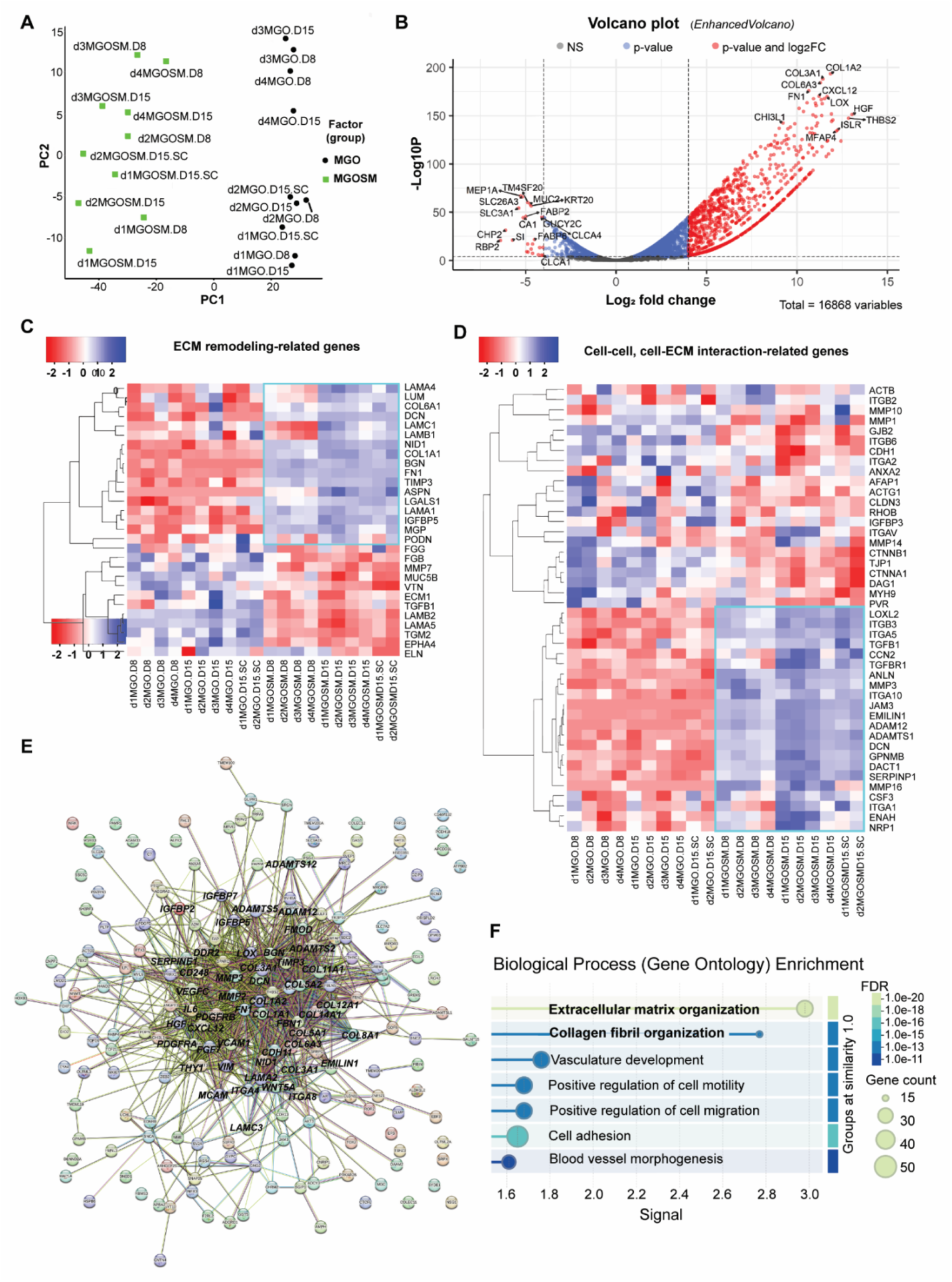
Transcriptomics reveal enhanced ECM remodeling by mesenchymal cells. (A) PCA graph shows the contribution of the top two principal components to the variance in mRNA expression between the organoid-only (O) and co-culture (OSM) conditions in Matrigel (MG). The relation between the top two principal components (PC1 and PC2) highlights the separation of the MGO and MGOSM samples. (B) Volcano plots showing log2 fold change (FC) versus significance (p-value) calculated for 16868 genes for the two independent populations, highlighting enriched genes for the MGO and MGOSM conditions. Dots in red indicate log2FC <-4 or >4, and with adjusted p-value <0.0001. Heatmaps were made out of overlapped genes between the 16868 genes and previously reported genes that are associated with ECM-remodeling (C), cell-cell & cell-ECM interaction (D). (E) Interaction networks of the top 200 most differently expressed genes between O and OSM. (F) Gene Ontology analysis reveals the top involved signaling pathways for the top 200 differently expressed genes between O and OSM.

**Figure 7.**
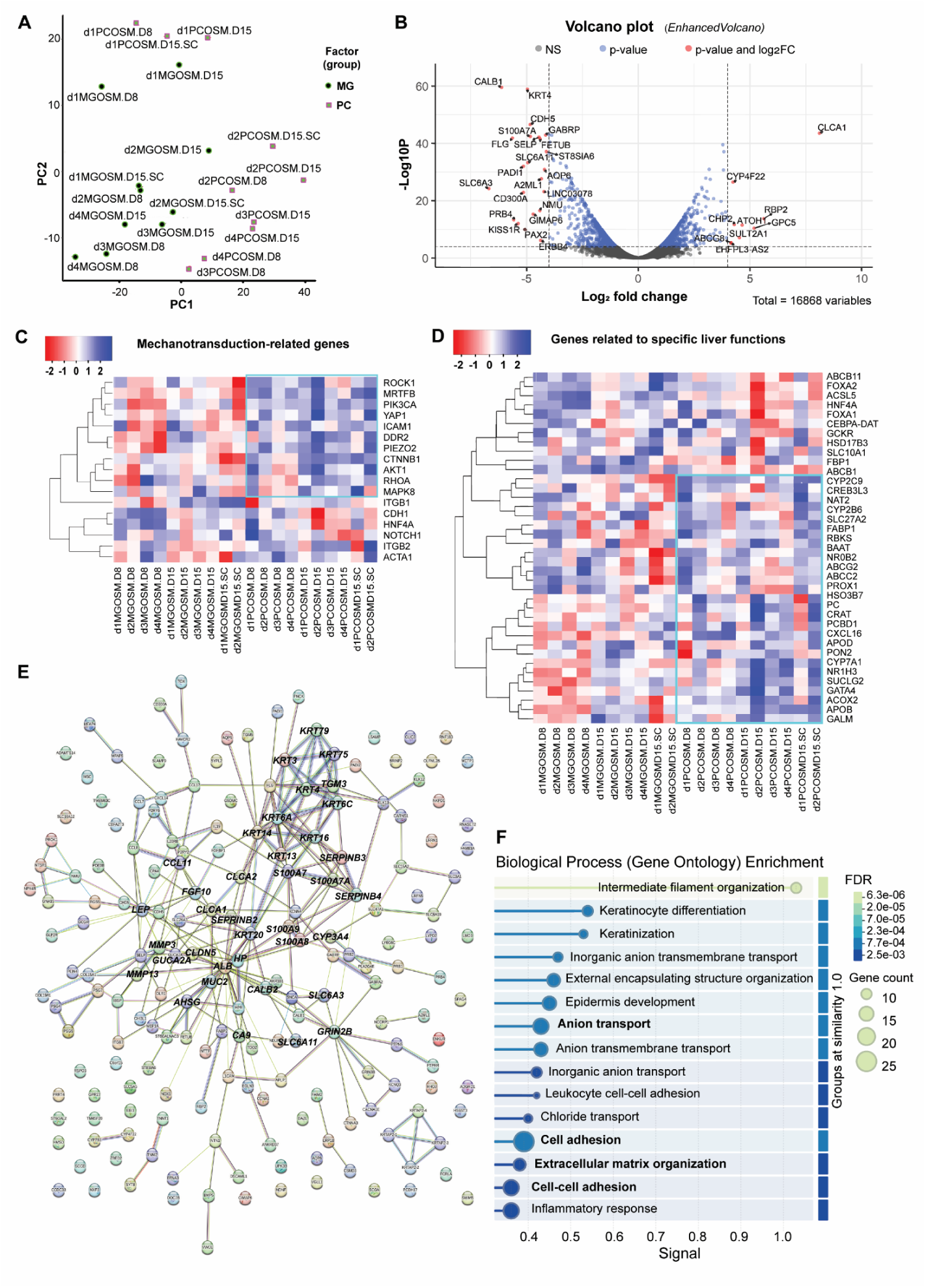
Transcriptomic comparison of BLTs in Matrigel and PC. (A) PCA graph shows the contribution of the top two principal components to the variance in mRNA expression between the Matrigel (MG) and PIC + Collagen (PC) conditions for co-cultures with OSM to form BLTs. The relation between the top two principal components (PC1 and PC2) highlights the separation of the MGOSM and PCOSM samples. (B) Volcano plots showing log2 fold change versus significance (p-value) calculated for all genes for the two independent populations, highlighting enriched genes for the MGOSM and PCOSM conditions. Dots in red indicate log2FC <-4 or >4, and with adjusted p-value <0.0001. (C-D) Heatmaps were made out of genes reported to play a role in mechanotransduction, or specific liver functions. (E) Interaction networks of the top 200 most differently expressed genes between MG and PC conditions. (F) GO analysis reveals the top involved signaling pathways for the top 200 differently expressed genes between MG and PC conditions.

### Transcriptomics Reveal Enhanced ECM Remodeling By Mesenchymal Cells

The PCA graph displays clear clusters of O (right) and OSM (left) conditions (**Figure 6A**). Differential expression analysis, shown in a volcano plot, reveals the upregulation of various ECM-related genes in OSM, especially multiple types of collagens, including *COL1A2*, *COL3A1*, *COL6A3* (**Figure 6B**). Importantly, hepatocyte growth factor (*HGF*) is among the highest upregulated genes, which could be an explanation for the faster organoid formation and growth in OSM than in O condition (Figure 1B). Compared to O condition, OSM expressed lower *SLC26A3*, *RBP2*, *FABP2*, *FABP6*, *MUC2*, *CLCA1*, *CLCA4*. Gene ontology (GO) analysis provides information on significant differences in biological processes between O and OSM. The involved activated processes include “collagen-containing extracellular matrix”, “extracellular matrix”, “extracellular matrix organization”. (**Figure S5A** left). While the suppressed biological processes are mainly related to the MHC class II protein complex, which may well result from mesenchymal cells (Figure S5A right). The gene set enrichment analysis (GSEA) shows that the representative GO signaling pathways, including “collagen-containing extracellular matrix”, “extracellular matrix”, “extracellular matrix organization” were all upregulated, while the MHC class II protein complex signal pathway was suppressed (**Figure S5B**). To have a closer look at the ECM remodeling and cell-cell and cell-ECM interaction-related gene expression, we applied the normalized gene counts to overlap with lists of previously reported genes (**Table S1**). The selected genes, displayed as heatmaps, show clear trends of higher or lower expressed genes in O or OSM **(Figure 6C, 6D**). For example, genes (*LAMA1, LAMB1, LAMC1, NID1, and COL1A1*) for the most abundant ECM components (laminin-entactin complex and collagen) are all highly expressed in OSM conditions (Figure 6C). In turn, several genes for cell-cell and cell-ECM interaction are also highly expressed in OSM conditions. On the contrary, the other genes highly expressed in O are lowly expressed in OSM (**Figure 6D**). In addition, we observed a similar pattern of gene expression between O and OSM for autocrine and paracrine effect-related, proliferation and apoptosis-related, and even for specific liver functions-related genes (**Figure S5C-E**). We then selected the top 200 most differentially expressed genes between O and OSM to focus on the most outstanding differences and related biological processes. The gene interaction shows that the core of these top 200 differentially expressed genes is mostly related to ECM remodeling, especially laminin-entactin, various collagens and MMPs. Other genes near the core of the network are related to “cell adhesion and growth” and “liver regeneration”, including *HGF*, *ITGA8*, *IL-6* (**Figure 6E**). GO analysis further verified that the enriched biological processes by these top 200 genes are “extracellular matrix organization”, “collagen fibril organization” and “cell adhesion”, which are important for ECM remodeling (**Figure 6F**). Taken together, the transcriptomic results reveal that mesenchymal cells enhanced ECM remodeling in BLTs.

### Transcriptomics Indicate That PC Promotes BLT Maturation Via Enhanced Mechanotransduction

We have previously found that PC enhanced the maturation of BLTs (Figure 4), but the underlying mechanism is not clear. Therefore, we compared the OSM-derived BLTs in MG and PC conditions via transcriptomic analysis. The PCA shows clustering of MG (left) and PC (right) conditions except for donor 1(d1)-PC, which is above d1-MG (**Figure 7A**). Upregulated genes in PC are related to metabolism, including *CYP4F22*, and *RBP2*, while inflammatory response-related genes, such as *S100A7A* and *CD300A,* are lower expressed in PC (**Figure 7B**). GO analysis revealed that multiple ribosomal subunit-related biological processes are activated in PC, indicating enhanced gene translation and active protein synthesis, including collagen trimer (**Figure S6A** left). Interestingly, several epidermis development-related processes are suppressed in PC (Figure S6A right). Graphs by GSEA show the representative GO signaling pathways, including the activated “Collagen trimer”, “cytosolic large ribosomal subunit”, “regulation of cardiac muscle cell membrane repolarization”, and the suppressed “transmembrane signaling receptor activity” (**Figure S6B**). The heatmap shows that many mechanotransduction-related genes are upregulated in PC compared to MG, including *YAP1, PIEZO2, DDR2, CTNNB1* (**Figure 7C**). Consistent with previous qPCR results, genes related to specific liver functions show an overall higher expression in PC than in MG. These highly expressed genes cover broad liver functions, including bile synthesis (*NR0B2, BAAT, CYP7A1, HSO3B7, ACOX2*) and bile transport (*ABCC2, ABCG2*), drug metabolism (*CYP2B6, CYP2C9, NAT2*), and metabolic activities related to glucose (*GALM, RBKS, PC, SUCLG2*), fat (*FABP1, CRAT, SLC27A2*), cholesterol (*APOB, APOD, PON2, CXCL16*) (**Figure 7D**). Interestingly, a distinct group of transcription factors are upregulated (*NR1H3, PCBD1, GATA4, CREB3L3*), whereas others are downregulated (*FOXA1, FOXA2, CEBPA-DAT, HNF4A*) in PC, which is opposite to the trend in MG (**Figure 7D**). As PC is a well-defined hydrogel, the pattern of cell-cell and cell-ECM interaction and ECM remodeling within BLTs with PC may well be different from that with MG. Of note, around half of cell-cell and cell-ECM interaction-related genes are higher expressed in PC while the other half are higher in MG, and the highly expressed genes in PC are further upregulated between day 8 (D8) and day 15 (D15) of differentiation, while the lower expressed genes in PC are downregulated between D8 and D15 (Figure S6C). This phenomenon indicates a dynamic change of gene expression with time; thus, we later conducted a separate comparison between D8 and D15, focusing on O and OSM, respectively. We further checked the genes associated with ECM remodeling and found that around one-third of these genes are higher expressed in PC and also upregulated from D8 to D15. These upregulated genes are closely related to the laminin-entactin complex (*LAMA1, LAMB1, LAMC1, NID1*) that we previously reported to be important for liver progenitor/stem cell growth (Figure S6D), which may suggest the maintenance of a stem cell niche in BLTs with mesenchymal cells in PC. On the other hand, genes highly expressed in MG are associated with chondrogenesis (*ASPN*), bone formation (*ECM1, BGN*), and one of the major ECM components *COL1A1* together with the *TIMP3* - an inhibitor of matrix metalloproteinases - (Figure S6D), which are not ideal for the maintenance of hepatic phenotypes. Next, we utilized the top 200 most differentially expressed genes between MG and PC to focus on the most outstanding differences and related biological processes. The gene interaction shows that there are two cores (*ALB* and keratin family) in the interaction network. Genes highly connected to *ALB* include *CYP3A4, GUCA2A*, and the keratin gene family core connects *KRT4, KRT13, KRT14, KRT3, KRT6A*, etc. (**Figure 7E**). The enriched biological processes by these 200 genes include “intermediate filament organization”, which shows the strongest signal, and most of the genes are associated with “cell adhesion”, “anion transport”, “cell-cell adhesion”, “inflammatory response”, and also “ECM organization” (**Figure 7F**). It is known that the main function of intermediate filaments is to provide support and structure for cells, so the strong intermediate filament organization indicates enhanced mechanical strength and resistance to shear stress for BLTs in PC. These results indicate that the enhanced mechanotransduction could be the reason for the promoted maturation of BLTs in PC.

### Transcriptomics reveal stronger influences of DS on OSM than on O

Clustering of static culture (SC) and dynamic suspension culture (DS) samples in the PCA graph is not clear due to tremendous differences in cellular composition and utilized hydrogels (**Figure S7A**). GO analysis displays that the most activated biological processes in the DS condition are associated with biosynthesis or metabolic processes besides “Extracellular matrix” (**Figure S7B**, left). Surprisingly, the most suppressed biological processes are related to epidermis development (**Figure S7B**, right). As the GO analysis focused on the comparison between SC and DS conditions, the influence of DS on O and OSM remains unknown. To investigate if there are different influences of DS on O and OSM, we made heatmaps as described. Most cell-cell and cell-ECM interaction-related genes are expressed higher in OSM than in O, and their expression levels are even higher under DS than under SC conditions (**Figure S7C**). Only the minority of genes associated with cell-cell and cell-ECM interaction are higher expressed in O than OSM, with more pronounced differential expression levels under DS conditions (**Figure S7C**). In terms of ECM remodeling-related gene expression, although there are clear clusters of genes highly expressed in O but lower in OSM, or genes highly expressed in OSM but lower in O, the expression levels between SC and DS are comparable within O and OSM conditions, respectively (**Figure S7D**). Of note, the overall expression of autocrine and paracrine effect-related genes is lower in OSM under DS conditions, while these genes retain similar expression levels in O under both SC and DS conditions (**Figure S7E**). As expected, mechanotransduction-related genes are overall highly expressed under DS conditions, especially higher in OSM than in O (**Figure S7F**). Taken together, the DS condition affected biosynthetic and metabolic processes and influenced cell-cell and cell-ECM interactions but not ECM remodeling. Also, DS condition suppressed the autocrine and paracrine effects (mainly contributed by mesenchymal cells) in BLTs. The overall influence of the DS condition on BLTs is stronger on OSM (co-culture) than on O (organoid-only).

### Long-term Functional Maintenance Relies On Both Cellular and Extracellular Stimuli

We found differentially expressed hepatic proteins between D8 to D15 (**Figure 5**), and transcriptomic comparison of different hydrogels (MG vs PC) and culture conditions (SC vs DS) also indicated changes of gene expression patterns from D8 to D15 (**Figure 6-7, S5-S6**) and between O and OSM (**Figure S7**). To investigate the major contributing factors for long-term maintenance of hepatic functions of BLTs, we further compared BLTs between D8 and D15 within O and OSM conditions, respectively. The PCA for O samples between D8 and D15 displays clear separation of samples by donors (d1, d2, d3, d4) with obvious trends of D8 samples located above D15 samples within each donor (**Figure S8-1A**). Instead, the PCA for OSM displays clustering of samples by hydrogels (PC on the right and MG on the left), in addition to donors (**Figure S8-2A**). GO analysis displays distinguishable activated and suppressed biological processes between O and OSM regarding their comparison between D8 and D15 conditions. Ribosome-related biological processes are activated on D15 compared to D8 for O conditions, indicating upregulated protein synthesis (**Figure S8-1B**, left). However, some biological processes related to hepatic functional maintenance are suppressed on D15 for O conditions, including “Digestion”, “Digestive tract morphogenesis”, and “Blood microparticle” (Figure S8-1B, right). Compared to OSM samples on D8, OSM samples on D15 show several suppressed biological processes that are filament-related or related to epithelium maintenance, with counts all less than 50 (**Figure S8-2B**, right). Of note, biological processes related to “Vascular/Blood vessel development” and “Tube morphogenesis” are highly activated (**Figure S8-2B**, left), which may contribute to vascularization and engraftment of the BLTs after *in vivo* transplantation.

We further investigated the genes related to “Cell-cell & cell-ECM interaction”, “ECM remodeling”, “Proliferation & apoptosis”, and “specific liver functions” as described above and displayed them as heatmaps. All heatmaps of O samples show no consistent trend of changes in gene expression between D8 and D15 (**Figure S8-1C-F**). Unlike O samples, OSM samples always show clear gene expression trends between D8 and D15, in all four groups of genes shown in the heatmaps. The majority of genes related to cell-cell & cell-ECM interaction are upregulated from D8 to D15, including *ITGA1, ITGAV, ITGB3, DCN, TGFB1*, and *TGFBR1*; the remaining genes of this group show lower expression levels on D15 than on D8, including *CDH1, ITGB2, CLDN3, PVR* (**Figure S8-2C**). Similar trends of gene expression changes are found in ECM remodeling-related genes, with higher expressed genes downregulated and lower expressed genes upregulated from D8 to D15 (**Figure S8-2D**). In particular, the upregulated genes include *LAMA1, LAMB1, LAMC1, NID1*, and *COL1A1*, and these genes encode the most abundant (>90%) ECM proteins, such as laminin-111, entactin, and collagen. It is known that the laminin-entactin complex is important for liver stem/progenitor cell growth, indicating the possibility of maintaining a stem cell niche in the BLTs, contributing to the long-term maintenance of hepatic functions. Although around two thirds of displayed liver function related genes are downregulated from D8 to D15, the higher expressed genes are associated with key liver functions, such as bile synthesis (*CYP7A1, ACOX2, HSD3B7*), cholesterol (*CXCL16, APOD, APOB, PON2*), glucose (*PC, GALM, SUCLG2*), transcription factors (*GATA4, NR1H3*) (**Figure S8-2E**). The downregulation of bile transport-related genes (*ABCB1, ABCB11, ABCC2, ABCG2, SLC10A1*) may be due to the polarity change of organoids in the co-cultures with mesenchymal cells, especially in PC. Moreover, all highly expressed genes related to proliferation and apoptosis are downregulated from D8 to D15, such as *EGFR, DKK1, MELK, TGFBR2, CCNA2*, and *MKI67*, and the other well-known proliferative marker *PCNA* is also shown to be downregulated (**Figure S8-2F**). These results suggest that BLTs are lowly proliferative and are protected from apoptosis till D15. Based on the results comparing D8 to D15 samples, we found that cellular (incorporation of epithelial and mesenchymal cells) and extracellular factors (dynamic ECM mimicry) could perform interactive contributions in the long-term structural and functional maintenance of BLTs.

## Discussion

In this study, we demonstrate a novel method to bioengineer liver tissue using a co-culture system of human liver organoids with hepatic stellate cells (HSCs) and mesenchymal stromal/stem cells (MSCs). Liver organoids used in this study are intrahepatic cholangiocyte organoids (ICOs), which consist of epithelial cells that are bipotential and can be differentiated into hepatocyte-like and cholangiocyte-like cells with a respective suitable differentiation medium (DM) (*6*, *28*, *29*). Hepatocytes and cholangiocytes are the parenchymal cells of the liver, making up ∼80% of the liver cell mass. Here, we combined epithelial organoids with mesenchymal cells (HSCs and MSCs) to induce spontaneous tissue formation and create hepatic tissues representing the cellular complexity of the liver. HSCs are liver mesenchymal cells, well-known for their important roles in liver regeneration (*9*, *30*). MSCs are frequently included in co-culture models to enhance hepatic function in *in vitro* models (*31*). Previously, Cordero-Espinoza *et al*. reported dynamic cell contacts between periportal mesenchyme and ductal epithelium in mouse liver regeneration and murine liver organoids (*32*). Recently, Haaker *et al.* published the results of enhanced proliferation of liver organoids when co-cultured with HSCs (*33*). However, to our knowledge, the individual influences of HSCs and MSCs on human organoid expansion, differentiation, and tissue formation remained unknown. In our study, we compared the morphological differences of different organoid cultures: organoids-only (O), organoids co-cultured with HSCs (OS), organoids co-cultured with MSCs (OM), and organoids co-cultured with HSCs and MSCs (OSM). The bright-field pictures showed that all co-cultures resulted in enhanced organoid formation and growth in organoid expansion medium (EM) (Figure 1B-D), which could be promoted by direct cell-cell contact together with soluble factors secreted by mesenchymal cells as reported previously (*32*, *33*). Our transcriptomic analyses indicate that ECM remodeling by mesenchymal cells could also contribute to the enhanced organoid formation and growth, especially since ECM components like laminin-entactin complex are produced (Figure 6). Once changed to organoid DM to induce hepatic function, all cultures showed condensed morphologies with obvious differences between organoid-only and co-cultures: in organoid-only conditions, individual organoids adapted a more condensed phenotype, while co-cultures not only became darker than organoid-only, but also spontaneously fused into more compact tissue-like structures, with few visible single mesenchymal cells (Figure 2B). This spontaneous tissue formation is consistent with the concept of mesenchymal cell-driven iPSC condensation into liver buds reported by Takebe *et al.* (*12*). However, there was no distinguishable morphological difference among OS, OM, and OSM. Therefore, we characterized different cultures on the mRNA level by qPCR assays. The results showed that in EM conditions, the only significant difference was the lower expression of stem cell marker *LGR5* in HSC-containing co-cultures compared to organoids-only (Figure 1E). In contrast, in DM conditions, the expression levels of most hepatocyte markers were lower in co-cultures containing HSCs, while the expression levels of *Ki67* were higher (Figure 2C). These results indicate that HSCs could have maintained the organoids in a proliferative state, preventing them from differentiating. However, we cannot exclude that the lower expression of hepatocyte markers (and the trend of downregulation of *LGR5*) could have resulted from the dilution by proliferative HSC mRNA/cDNA. In contrast to the ambiguous role of HSCs in bioengineered liver tissues (BLTs), we saw that MSCs played a relatively clear role in the co-cultures by enhancing organoid formation and growth in EM, promoting spontaneous tissue formation, and supporting hepatic function. Although the exact function HSCs performed in BLTs was not clear, we continued this study with the tri-culture (OSM) condition to mimic the cellular complexity and multicellular cooperation of the *in vivo* situation (*9*, *34*). For future studies, it would be interesting to also include Kupffer cells and liver sinusoidal endothelial cells to better mimic the immunological functions of the native liver (*35*). Besides mesenchymal cells, Kupffer cells and endothelial cells make up most (∼70%) of the non-parenchymal cells in the liver, and as resident antigen-presenting cells, they are crucial in maintaining tolerance under noninflammatory conditions (*36*, *37*).

Besides cellular factors, we showed that the culture method played a vital role in tissue formation. After around 7 days of static culture (SC), we observed that some cells had migrated to the bottom of Matrigel droplets, where they attached and proliferated on the plastic wells, which may have contributed to a higher expression of proliferation markers in the co-cultures during differentiation. To avoid this, we utilized low-attachment plates and placed them on a horizontal shaker as a dynamic suspension culture (DS) method. Remarkably, the DS method not only avoided cell attachment but also promoted spontaneous tissue formation and even enhanced the maturation of BLTs, including the ECM markers *FN* and *VTN* (Figure 3C-D). The higher *FN* expression in co-cultures probably indicates the potential ECM remodeling by mesenchymal cells, and the trend of upregulated expression of *VTN* in DS is consistent with other hepatocyte markers. This enhanced maturation may be mainly induced by fluidic shear forces caused by the DS condition. Previously, we reported that suspension culture of liver organoids in spinner flasks induced better differentiation compared to SC (*8*), and Takahashi *et al.* developed a protocol for the differentiation and maturation of human intestinal organoids completely in suspension (*26*). All those studies in suspension underline the importance of a dynamic (micro-) environment including shear stress for tissue formation and regeneration. Our transcriptomic results suggest that the enhanced maturation was not only induced by mechanotransduction changes in the DS condition, but also influenced by “cell-cell and cell-ECM interaction” and “autocrine and paracrine effects” (Figure S7). In future tissue engineering approaches, continuous fluid flow by perfusion might induce even better tissue fusion and maturation than the DS system used in this study.

Using hydrogels as ECM mimicry not only provides cells with a 3D microenvironment, it can also offer the possibility to adjust mechanical properties and biological functions according to the needs of a particular tissue (*38*, *39*). Although the DS method upregulated the expression of hepatocyte markers on the mRNA level, the key metabolic protein CYP3A4 was hardly detected in Matrigel cultures (Figure 3E). Then, we replaced Matrigel with PIC-Collagen-I as a well-defined hydrogel. Previously, PIC hydrogels were used for the culture of human liver organoids and mammary gland organoids (*16*, *17*) and for the differentiation of cholangiocyte organoids (*40*). Collagen-I has been widely applied as a biomaterial in TERM (*41*), and primary mouse hepatocytes and human HepaRG cells secreted higher levels of urea and albumin in collagen-I microspheres than in most other tested ECM components (*27*). In our study, the co-cultures benefited from the combination of the synthetic thermoreversible PIC supplemented with collagen I (PC). PC enabled organoid co-cultures with MSCs and HSCs to spontaneously form compact liver tissues (Figure 4B) and significantly induced hepatic marker expression, including the expression of CYP3A4 on the protein level (Figure 4C, 4E, 7E). Our transcriptomic results suggest that PC could have promoted BLT maturation via enhanced mechanotransduction (Figure 7C-D). Although the expression of some hepatic markers showed a downregulation during long-term culture, this is more likely due to the continuous utilization of the DM that is developed for liver organoid differentiation but not ideal for the maintenance of hepatic functions. Importantly, BLTs formed in PC retained a vital portion of hepatic functional markers during long-term maintenance of BLTs (Figure S8-2E). Moreover, PC could have created a stem cell niche with abundant laminin-entactin complex via ECM remodeling and cell-cell and cell-ECM interactions, which may contribute to the long-term maintenance of hepatic functions (Figure S8-2C-D). Furthermore, since both PIC and collagen-I are thermoresponsive, the hydrogels can easily be removed after liver tissue formation, resulting in xeno-free BLTs suitable for *in vivo* applications.

In conclusion, we have established a straightforward method to bioengineer human liver tissues *in vitro* by co-culturing human liver epithelial organoids (ICOs) with mesenchymal cells (HSCs and MSCs) embedded in a well-defined thermoreversible PC hydrogel on an orbital shaker. This study incorporated several parameters to investigate the interactive effects among different influencing conditions for tissue engineering, using the liver as a showcase: A. Spontaneous multicellular organization in co-culture conditions. B. Dynamic stimuli induced by suspension culture on a shaker. C. Well-defined hydrogels as ECM mimicry. D. Long-term maintenance of hepatic functions. The BLTs provide valuable models for toxicity tests, drug selection, disease modelling, and lay the foundation for bioengineering larger tissues, promising for *in vivo* transplantations. The tissue engineering strategy described here paves the way for further development of additional biomimetic approaches not only for the liver but also for other tissue types.

## Materials and Methods

### Experimental Design

The experimental design of this study is shown in **Figure 8**.

**Figure 8.**
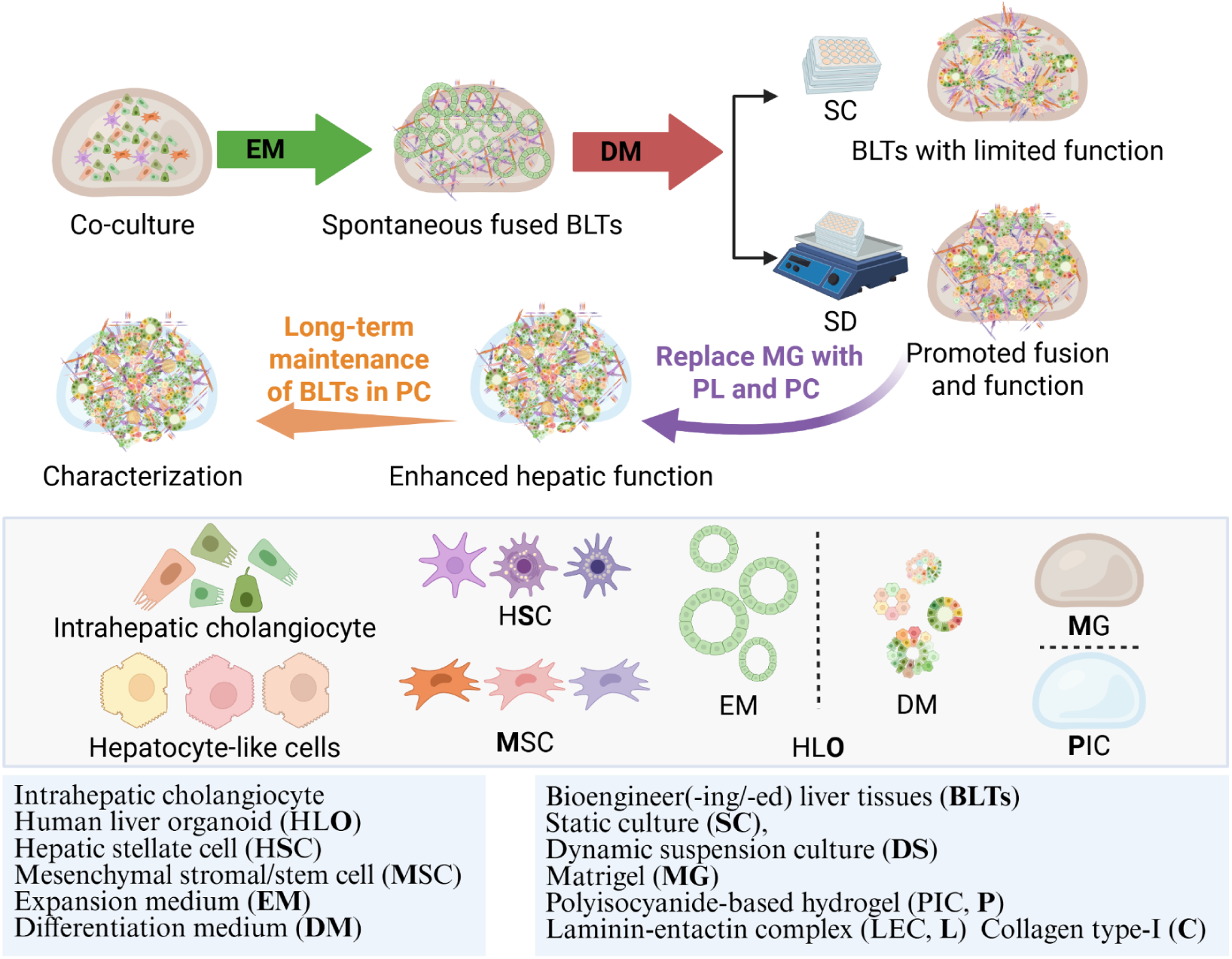
The schematic research line of experimental design for LTE.

### Medium preparation

Advanced DMEM/F12 (Gibco) supplemented with 1% (v/v) penicillin-streptomycin (Gibco), 1% (v/v) GlutaMax (Gibco), 10 mM HEPES (Gibco), was used as the basal medium to make organoid initiation medium, expansion medium (EM), differentiation medium (DM), and to wash organoids during passaging or sample collection.

Other major materials used for organoid isolation and maintenance are as follows: type II collagenase (Gibco), dispase (Gibco), Matrigel^TM^ (Corning), and non-attaching 24-well plates (M9312, Greiner, Merck). Certified Fetal Bovine Serum (FBS) was ordered from Gibco (16000–044). Organoid initiation medium was prepared by using 70% EM and adding 30% Wnt-conditioned medium (homemade).

EM was made based on basal medium, supplemented with 2% (v/v) B27 supplement without vitamin A (Invitrogen), 1% (v/v) N2 supplement (Invitrogen), 10 mM nicotinamide (Sigma-Aldrich,), 1.25 mM N-acetylcysteine (Sigma-Aldrich), 10% (v/v) R-spondin-1 conditioned medium (the Rspo1-Fc-expressing cell line was a kind gift from Calvin J. Kuo), 10 µM forskolin (FSK, Sigma-Aldrich), 5 µM A83-01 (transforming growth factor b inhibitor; Tocris Bioscience), 50 ng/mL EGF (Invitrogen), 25 ng/mL HGF (Peprotech), 0.1 µg/mL FGF10 (Peprotech) and 10 nM recombinant human (Leu15)-gastrin I (Sigma-Aldrich).

DM was also based on basal medium, supplemented with 1.25 mM N-acetylcysteine (Sigma-Aldrich), 2% (v/v) B27 supplement without vitamin A (Invitrogen), 1% N2 supplement (Invitrogen), 50 ng/mL EGF (Invitrogen), 10 nM recombinant human (Leu15)-gastrin I (Sigma-Aldrich), 25 ng/mL HGF (Peprotech), 100 ng/mL FGF19 (Peprotech), 500 nM A83-01 (Tocris Bioscience), 10 µM DAPT (Selleckchem), 25 ng/mL BMP-7 (Peprotech), and 30 µM dexamethasone (Sigma-Aldrich). DM was changed every 2–3 days for around 8 days.

Hepatic stellate cell growth medium (HSCGM) was made from Dulbecco’s modified Eagle’s medium (DMEM; Thermo Fisher Scientific), 10% (v/v) FBS (Gibco), and 100 U penicillin/streptomycin (Gibco).

Mesenchymal stromal /stem cell growth medium (MSCGM) was based on DMEM, low glucose, pyruvate (Gibco), supplemented with 10% (v/v) FBS (Gibco), and 100 U penicillin/streptomycin (Gibco).

### Organoid isolation

Human liver organoids used in this study were intrahepatic cholangiocyte organoids (ICOs) (*42*). Human liver tissues were obtained during liver transplantation at the Erasmus Medical Center Rotterdam, in accordance with the ethical standards of the Helsinki Declaration of 1975. Using the tissue for research purposes was approved by the Medical Ethical Council of the Erasmus Medical Center and informed consent was provided by patients (MEC-2014-060). ICO lines were established and cultured as previously described (*6*). Briefly, to establish organoid lines, liver tissues were cut into small pieces, followed by enzymatic digestion with 0.125 mg/mL type II collagenase and 0.125 mg/mL dispase in basal medium containing 1% (v/v) FBS. The supernatant was collected every hour. The procedure of tissue digestion and the supernatant collection was repeated three times. Collected cells were washed in basal medium containing 1% FBS and centrifuged at 400 x *g* for 5 min. The cells were resuspended in Matrigel at a concentration of ∼500 cells per μL, then seeded as droplets in non-attaching 24-well plates. Organoid initiation medium was added after approximately 15 min incubation at 37°C, 5% CO_2_ in air.

### Cell passaging

For organoid expansion, fragments or single cells from organoids were plated within Matrigel droplets, and EM medium was added after gelation. The medium was changed every 2–3 days. Organoids were passaged by mechanical disruption once a week or by single cell dissociation every 10 days, both at an average split rate of 1:3-6 depending on density. In detail, when liver organoids were almost confluent in Matrigel droplets, they were either passaged as fragments or single cells. For passaging as fragments, organoids were collected in 15 mL Eppendorf tubes containing cold basal medium. Then, the organoids were centrifuged at 4°C at 400 x g for 5 min. After removing the supernatant, around 200 µL of cold basal medium was added to the tube and organoids were disrupted mechanically by pipetting up and down until they were small clusters. For passaging with single cells the medium was removed, and 1 mL of Tryple-Express was added to one well of a 12-well plate; then, the Matrigel droplets were disrupted mechanically by pipetting up and down for 7-10 times with 1 mL Eppendorf tips, and the plate was incubated at 37°C for around 30-60 min until the organoids became almost single cells. After that, the single cells or fragments were collected in 15 mL tubes, and the tubes were filled with cold basal medium (for enzymatic passaging with Tryple-Express, the basal medium contained 3-5% (v/v) FBS) and centrifuged again to get a pellet of organoid cells. Once the supernatant was removed, the organoid cells were resuspended in Matrigel and plated in a 24-well plate (50 µL/well). EM was added (500 µL/well) after approximately 15 min incubation at 37°C. All cultures were kept in a humidified atmosphere of 95% air and 5% CO_2_ at 37°C.

For HSCs and MSCs passaging, a 1:4–1:8 split ratio was performed once every 5–10 days. More specifically, once the cultures were almost confluent, cells were surpassed by using Tryple-Express (Gibco) for 5 min at 37°C. Cells were collected and centrifuged at 400 x *g* for 5 min at 4°C. After centrifugation, the supernatants were removed, and the cells were cultured in T75 flasks with 10 mL of HSCGM or MSCGM for HSCs and MSCs, respectively. All cultures were kept in a humidified atmosphere of 95% air and 5% CO_2_ at 37°C.

### Cell seeding and Hydrogel preparation

Matrigel (Corning®, 356231), PIC (1k-PIC-P, NCN01, Noviogel), and Laminin-entactin complex (LEC, Corning) were thawed at 4°C 1-2 hours before use. The preparation of single cells from organoids, HSCs, and MSCs was described above. Once the single cells were ready for use after cell counting, the required amounts of cell suspensions were added to 1.5 mL Eppendorf tubes. Then, the cell suspensions were centrifuged at 400 x *g* for 5 min at 4°C to get cell pellets for later use. During the waiting steps, the Collagen Type I (collagen-1, Sigma-Aldrich, 08-115) was diluted to a certain concentration so that a fixed volume could be added to each mixture of cells and hydrogels. The final concentrations of Matrigel, PIC, LEC, and collagen-1 were as follows: Matrigel (MG, 10 mg/mL), PIC (1 mg/mL), LEC (3 mg/mL), and collagen-1 (1.5 mg/mL). The cell concentrations of each type of cell in different hydrogels were kept consistent: 1,200 organoid cells/µL, 200 HSCs/µL, and 200 MSCs/µL, which means the ratio of different types of cells was: Organoid: HSC: MSC= 1,200: 200: 200 or 6: 1: 1. There were in total six different combinations of cultures: organoid only (O), organoid with HSCs (OS), organoid with MSCs (OM), organoid with HSCs and MSCs (OSM), HSCs only (S), and MSCs only (M).

After mixing the cells with hydrogel solutions on ice, the mixtures were seeded in droplets in prewarmed 24-well plates (4-7 droplets/50 µL/well). Plates were incubated at 37°C for around 15 min for the gels to solidify. Once the hydrogels became solid, EM was added (500 µL/well). Then, EM was refreshed every 2-3 days for 10-11 days. All cultures were kept in a humidified atmosphere of 95% air and 5% CO_2_ at 37°C.

### Differentiation

To induce hepatic functions, liver organoids and their co-cultures were primed for 3 days in EM with the addition of 25 ng/mL BMP-7 (Gibco), after which the medium was changed to DM. DM was changed every 2–3 days for 8 days or 14 days (for long-term assays).

### Static culture and dynamic suspension culture

For expansion, both the static culture (SC) and dynamic suspension culture (DS) conditions were cultured in a standard culture stove in respectively standard and low-attachment plates (Greiner). After expansion, the DS condition was placed on a horizontal shaker at a speed of 70 rpm for differentiation. Both conditions were kept in a humidified atmosphere of 95% air and 5% CO_2_ at 37°C.

### RNA isolation, cDNA synthesis, and qPCR assays

Trizol Reagent (Ambion, by Life Technologies, 15596018) was used to isolate RNA from organoids and other samples following the manufacturer’s instructions. Firstly, 0.5 mL of Trizol was added to each well (24-well plate) of organoids. After pipetting up and down for a couple of times, the cell suspensions were transferred into 1.5 mL RNase-free Eppendorf tubes. RNA quality and quantity were measured with DS-11 Spectrophotometer (DeNovix). Complementary DNA (cDNA) was synthesized with the iScript™ cDNA synthesis kit (Bio-Rad, 1708891) following the manufacturer’s instructions. Quantitative real-time PCR (qPCR) was used to determine relative expression of target genes using validated primers (Table 1) using the SYBR Green method (iQ SYBR Green Supermix, Bio-Rad). Normalization was carried out using reference genes *GAPDH* and ribosomal protein L19 (*RPL19*).

**Table 1.**
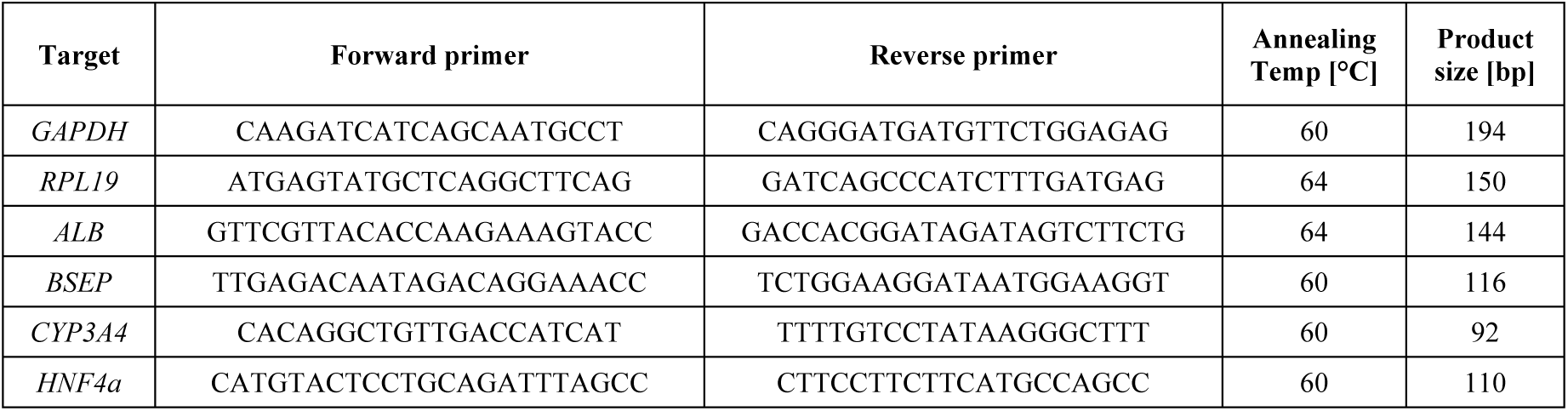
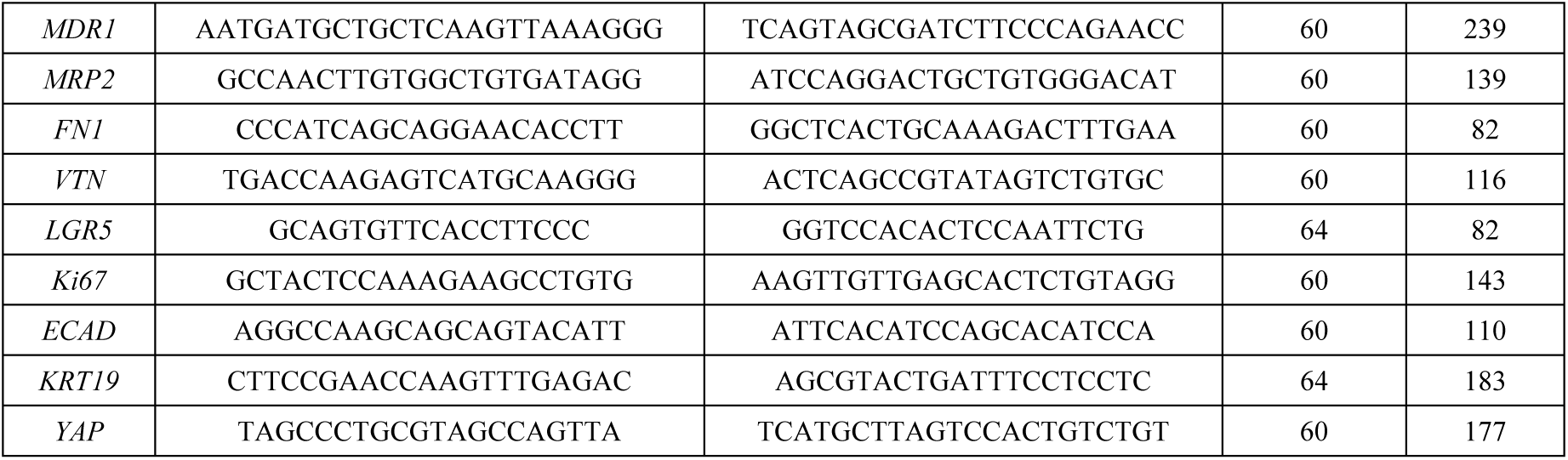
List of primers for gene expression profiling.

### Transcriptomic analysis

For whole transcriptome sequencing, RNA was isolated from organoids and other samples using the Trizol Reagent according to the manufacturer’s instructions. A minimum of 60 µL of RNA with RNA Integrity Numbers (RIN) ≥ 7, at a concentration of at least 10 ng/µL, was submitted for mRNA library preparation using poly(A) enrichment and sequenced at the Utrecht Sequencing Facility, following their internal protocols. Libraries were sequenced on the Illumina NextSeq2000 platform, generating paired-end reads (2 × 50 bp) with approximately 20 million reads per sample and 1 billion reads per flowcell. The mRNA reads were aligned to the human reference genome using STAR (version 2.4.2a, https://github.com/UMCUGenetics/RNASeq-NF/). Datasets were normalized for library size using the trimmed mean of M values (TMM). Fold changes were calculated with edgeR (*43*), and gene expression comparisons employed the exact test to compute p-values, adjusted for a 5% false discovery rate (FDR) using the Benjamini-Hochberg method. Genes with adjusted p-values < 0.05 were deemed differentially expressed. Principal component analysis (PCA) plots and heatmaps of differentially expressed genes were generated using ggplot2 (*44*) (https://cran.r-project.org/web/packages/ggplot2/citation.html]). Volcano plots were produced with EnhancedVulcano (*45*). (EnhancedVolcano: Publication-ready volcano plots with enhanced colouring and labeling. R package version 1.24.0, https://github.com/kevinblighe/EnhancedVolcano.), applying thresholds of P = 0.0001 and fold change (FC) = 4. Gene Set Enrichment Analysis (GSEA) (*46*)was conducted to identify enriched Gene Ontology (GO) terms among the differentially expressed genes.

### Immunofluorescent and histological staining

For immunofluorescent (IF), H&E and Periodic acid-Schiff (PAS) staining, organoids and bioengineered liver tissues (BLTs) were fixed with 4% (v/v) neutral buffered formalin containing 0.1% eosin and incubated at 37°C overnight. Fixed samples were dehydrated and embedded in paraffin or stored in 70% (v/v) ethanol at 4°C for up to 1 month; 4 μm thick paraffin sections were prepared for IF, H&E, and PAS staining. PAS staining and H&E staining were conducted as previously described (*8*) to evaluate the function of BLTs for glycogen storage and to compare the histology, respectively. To start the IF staining procedure, the paraffin sections were first heated at 62°C for 10 min and dewaxed by xylene, followed by rehydration in gradient ethanol concentrations from 100% to 70%. Then, sample sections were incubated in antigen retrieval solution for 30 min at 98°C. After balancing to room temperature, sample sections were treated with NH_4_Cl solution (20 mM) for 10 min to reduce background autofluorescence and blocked with 10% (v/v) goat serum for 1 h to avoid non-specific antibody binding. Next, primary antibodies against Albumin (ALB), Cytochrome P450 Family 3 Subfamily A Member 4 (CYP3A4), E-cadherin (ECAD), and Keratin 19 (K19) were added to the sections and incubated overnight at 4°C. After being washed with PBS containing 0.1% (v/v) Tween 20 for three times, sample sections were incubated with secondary antibodies (5 μM), including mouse anti-rabbit Alexa Fluor® 488, mouse anti-rabbit Alexa Fluor647, rabbit anti-mouse Alexa Fluor488 and rabbit anti-mouse Alexa Fluor647 (all from Molecular Probes). Nuclei were stained with DAPI (0.5 µg/mL, Sigma Aldrich). Detailed information on applied antigen retrieval methods, antibodies, dilutions, and incubation times are listed in **Table 2**.

**Table 2.** List of antibodies used.

| Primary antibodies diluted in antibody diluent (Dako) for IF |  |  |  |  |  |
| --- | --- | --- | --- | --- | --- |
| Antigen | Source and cat. number | Raised in | Dilution | Antigen retrieval | Incubation |
| ALB | Sigma, A6684 | Mouse | 1:1,000 | TE buffer for 30 min at 98°C | O/N at 4°C |
| CYP3A4 | Abcam, ab124921 | Rabbit | 1:100 |  |  |
| E-cadherin | BD Bioscience, 610181 | Mouse | 1:100 |  |  |
| KRT19 | Abcam, ab76539 | Rabbit | 1:150 |  |  |
| BSEP | Invitrogen, PA5-78690 | Rabbit | 1:500 |  |  |
| KRT18 | Santa Cruz, sc-51582 | Mouse | 1:100 |  |  |
| MRP3 | Abcam, ab3375 | Mouse | 1:100 |  |  |
| Secondary antibodies diluted in antibody diluent (Dako) |  |  |  |  |  |
| Antigen | Source and cat. number | Raised in | Dilution | Incubation |  |
| Anti-mouse | ThermoFisher A-11029 | Goat | 1:200 | 1 h at RT |  |
| Anti-rabbit Alexa | ThermoFisher A-11034 |  |  |  |  |
| Anti-mouse | ThermoFisher A-11004 |  |  |  |  |
| Anti-rabbit Alexa | ThermoFisher A-11036 |  |  |  |  |

### TEM analysis

Organoid-only (O-) and OSM-derived BLTs in Matrigel and PIC+collagen I (PC) hydrogels were fixed in half-strength Karnovsky fixative (2.5% Glutaraldehyde (EMS) + 2% Formaldehyde (Sigma)) in pH 7.4 0.1M PHEM buffer at RT for 2 hours. Fixed samples were rinsed and stored in 1% FA at 4°C until further processing. Post-fixation was performed with 1% OsO4, 1.5% K3Fe(III)(CN)6 in 1 M phosphate buffer pH 7.4 for 2 hours. BLTs were then dehydrated in a series of acetone (70% overnight, 90% 15 min, 96% 15 min, 100% 3x 30 min), and embedded in Epon (SERVA). 70nm ultrathin sections were cut (Leica Ultracut UCT), collected on formvar and carbon-coated transmission electron microscopy (TEM) grids, and stained with uranyl acetate and lead citrate (Leica AC20). Micrographs were collected on a JEM1011 (JEOL) equipped with a Veleta 2 k × 2 k CCD camera (EMSIS, Munster, Germany) or on a Tecnai12 (FEI Thermo Fisher) equipped with a Veleta 2 k × 2 k CCD camera (EMSIS, Munster, Germany) and operating SerialEM software. The alignment and montage of the stitched images were established in the IMOD software (https://bio3d.colorado.edu/imod/). The images were visualized and analyzed in FiJi (*47*)

### Microscopy and images analysis

Imaging of the cultures was performed using an EVOS FL Cell Imaging System (Life Technologies). Bright-field images were taken to track organoid morphology throughout expansion in different cellular formulations. Images were also taken to compare the morphology of organoids and their co-cultures, in EM and DM, embedded in Matrigel or PIC hydrogels, under SC or dynamic suspension culture (DS) conditions.

IF, H&E and PAS stainings were imaged with an Olympus BX51 microscope in combination with an Olympus DP73 camera.

### Intracellular and medium protein measurement

To quantify the intracellular and medium levels of ALB, ALAT, ASAT, and GLDH, BLTs were differentiated in PC or Matrigel droplets for 8 or 15 days as described above. BLTs were provided with fresh DM 24 h before being lysed in MilliQ water. ALB, ALAT, ASAT, and GLDH were measured in the cell lysates and for ASAT, also the medium supernatant, using a DxC-600 Beckman chemistry analyzer (Beckman Coulter). Values were normalized to live cell numbers.

### Ammonium Elimination Assay

For ammonium elimination assays, both organoid-only (O-) and OSM-derived BLTs were differentiated in PC or Matrigel droplets for 15 days as described above. BLTs were incubated with DM supplemented with NH_4_Cl (2 mm) for 24 h on both D8 and D15. After 24 h, D8 and D15 media samples were harvested and stored at −20 °C, respectively. Afterward, Tryple-Express (Gibco) was added to each well, and BLTs were trypsinized for cell counting. Cell counts were carried out using the TC20 automated cell counter (Bio-Rad). Viable cells were determined using the trypan blue exclusion assay. Ammonium concentrations were measured with the urea/ammonia Assay Kit (Megazyme). As a control, DM containing NH_4_Cl (2 mm) was incubated for 24 h without cells. Ammonia elimination rates were normalized to live cell numbers.

### Metabolic activity analysis with LC-MS/MS

The drug-metabolizing activity of BLTs was evaluated using a cocktail of parental compounds, as previously described (*48*). These included 5 μM Midazolam (BUFA; metabolites: 1’-Hydroxymidazolam, Sigma-Aldrich; and 1’-Hydroxymidazolam glucuronide, LGC Standards), 15 μM Phenacetine (Sigma-Aldrich; metabolite: Acetaminophen, Sigma-Aldrich), and 12 μM 7-Hydroxycoumarin (Umbelliferone, Sigma-Aldrich; metabolite: 7-Hydroxycoumarin glucuronide, LGC Standards). Cultures were exposed to the cocktail in DM in a humidified atmosphere at 37°C and 5% CO2 for 24 hours. 200 μL culture medium was collected in the 1.5-mL Short Thread Vials with the ND9 Short Thread Screw Caps (BGB Analytik), diluted 8 times with ultrapure MeOH, vortexed and stored at - 20°C until the quantification of the parental compounds and metabolites using LC-MS/MS, as previously reported (*48*). Briefly, prior to the analysis, samples were centrifuged for 10 min at 1500 g to precipitate any protein. Standards of the parental compounds and metabolites were prepared in the same medium (matrix) as samples. 1 μL solution was injected for measurements. Standards and samples were analyzed in a single run using a Shimadzu triple-quadrupole LCMS 8050 system with two Nexera XR LC-20AD pumps, a Nexera XR SIL-20AC autosampler, a CTO-20AC column oven and an FCV-20AH2 valve unit (Shimadzu). The compounds were separated on a Synergi Polar-RP column (150 × 2.0 mm, 4 μm, 80 Å) with a 4 × 2 mm C18 guard column (4 × 2 mm, Phenomenex). The mobile phase consisted of 0.1% (*v*/*v*) formic acid in ultrapure water (A) and 0.1% (*v*/*v*) formic acid in MeOH (B), and was set as 100% A (0-1 min), 100% to 5% A (1-8 min), 5% A (8-9 min), 5% to 100% A (9-9.5 min) and 100% A (9.5-12.5 min). The total run time was 12.5 min, and the flow rate was 0.2 mL/min. Peaks were integrated using the LabSolutions software (Shimadzu). For each compound, a limit of quantification (LOQ) was determined based on a standard curve and a limit of detection (LOD) was determined based on a noise signal. For each sample set, a dose (time 0 hours) and a blank (negative control) were measured. Amounts of metabolites were normalized to live cell numbers.

### Statistical analyses

Gene expression and functional analysis results were analyzed in the Excel sheets (Microsoft) and then transformed into graphs using GraphPad Prism 9. Tukey’s multiple comparisons test by One-Way or Two-Way ANOVA Multiple comparisons. The p-values (significance set at 0.05) are indicated in the respective figures. All average values described in the text and figures are mean ± SD (standard deviation).

## Acknowledgments

We thank Dr. Adam Myszczyszyn for his technical advice and assistance with the metabolic analyses, Cecila de Heus for technical assistance with transmission electron microscopy and Dr. Maya W. Haaker for kindly sharing hepatic stellate cells.

## Funding

Dutch Research Council NWO VENI (016.Veni.198.021) (KS)

China Scholarship Council (CSC201808310180) (SY)

China Scholarship Council (CSC201808620130) (ZW)

## Author contributions

Conceptualization: SY

Methodology: SY, ZW, NL, FvS

Investigation: SY, ZW, KS

Visualization: SY, ZW, NL, FvS

Supervision: KS

Writing—original draft: SY

Writing—review & editing: SY, KS, NL, FvS, ZW, BS, JM, LL

Funding acquisition: KS, SY, ZW

## Competing interests

The authors declare no competing interests.

## Supplementary Figures and Tables

**Figure S1.**
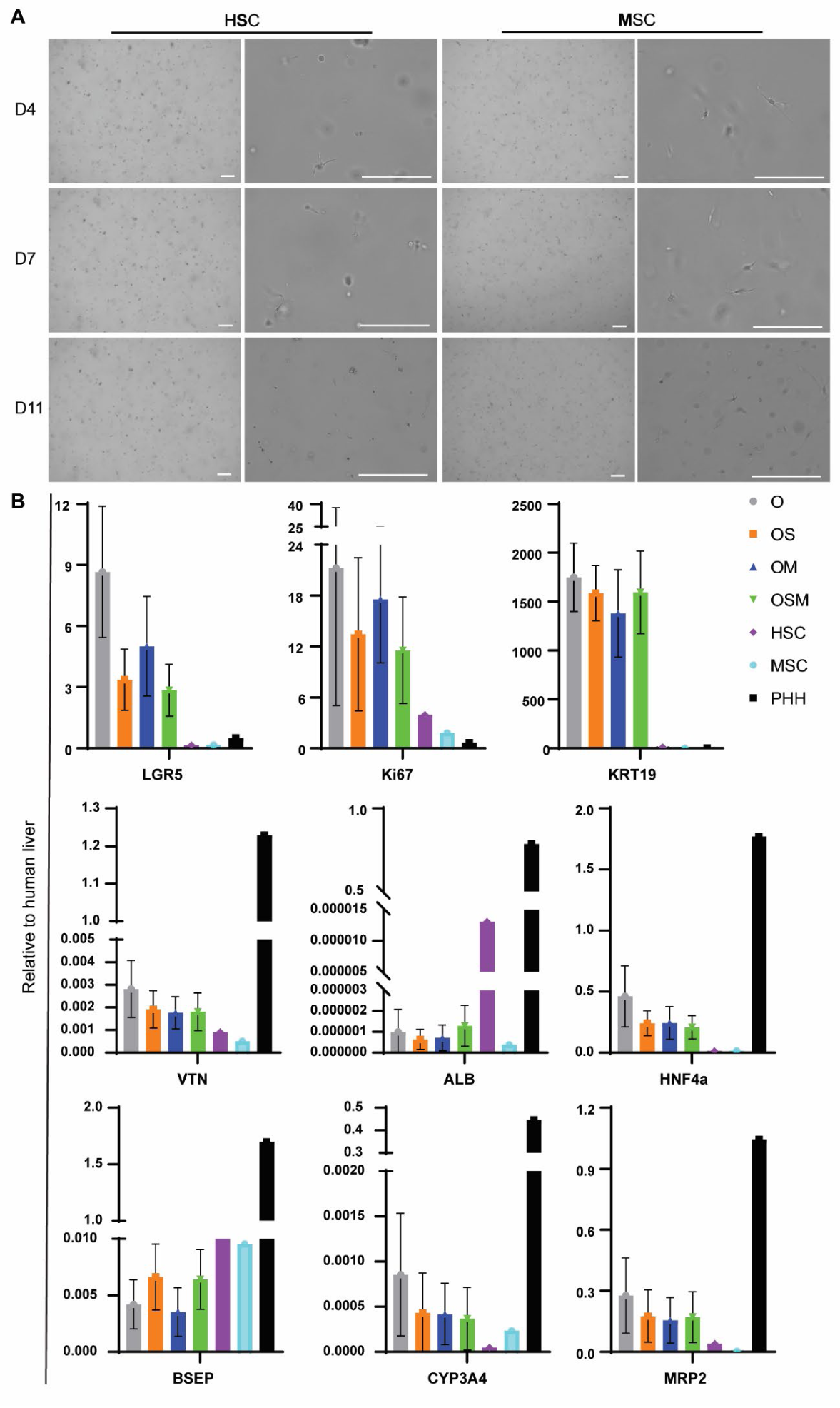
Expansion of mesenchymal cells in Matrigel. n=1. (A) Bright-field pictures showing the morphology of mesenchymal cells (HSC & MSC) in expansion medium (EM) at three time points, D4 (top), D7 (middle) and D11 (bottom). (B) Gene expression of mesenchymal cells only (HSC or MSC) expanded in Matrigel for 11 days, compared to primary human liver tissues. Stem cell/progenitor marker LGR5, proliferative marker Ki67, ductal marker KRT19, and hepatocyte markers HNF4a, ALB, CYP3A4, BSEP, MRP2, and VTN were used for the qPCR assays.

**Figure S2.**
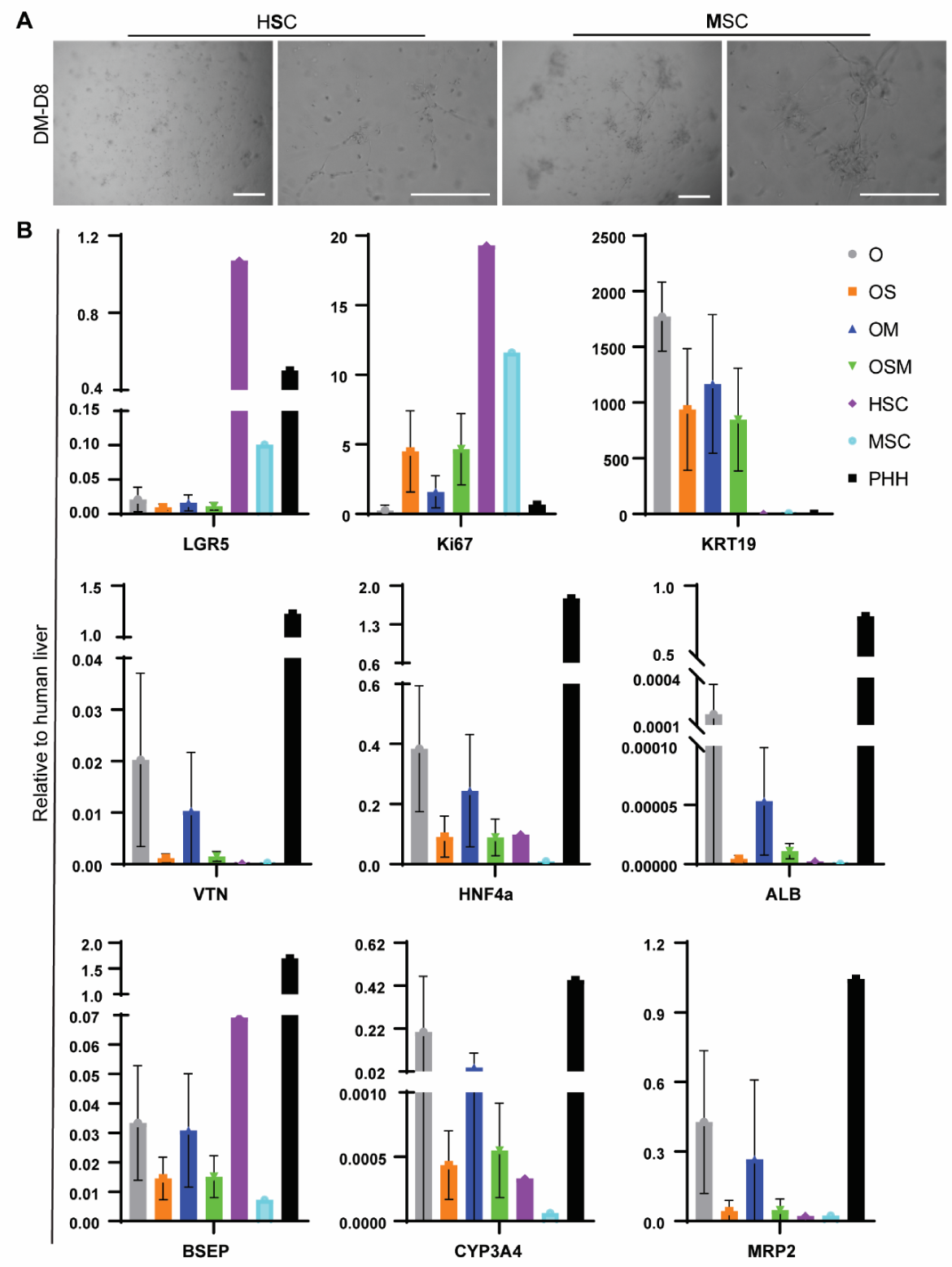
Mesenchymal cells in differentiation medium in Matrigel. n=1. (A) Bright-field pictures showing the morphology of mesenchymal cells (HSC & MSC) in differentiation medium (DM) for 8 days. (B) Gene expression of mesenchymal cells only (HSC or MSC) in Matrigel for 8 days, compared to primary human liver tissues. Stem cell/progenitor marker LGR5, proliferative marker Ki67, ductal marker KRT19, and hepatocyte markers HNF4a, ALB, CYP3A4, BSEP, MRP2, and VTN were used for the qPCR assays.

**Figure S3.**
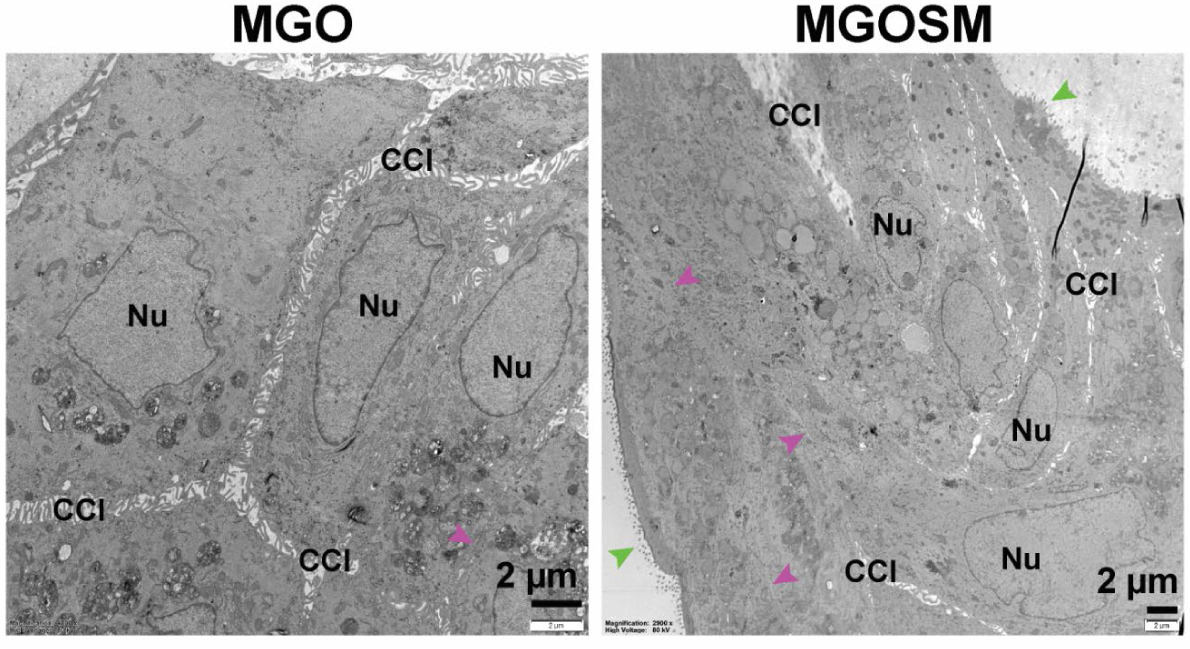
BLTs in MG obtained fewer structural characteristics than in PC. Subcellular characterization of BLTs in Matrigel (MG) with TEM analysis. Nucleus (Nu), tight junction (purple arrow), microvilli (MV, green arrow). n=4

**Figure S4.**
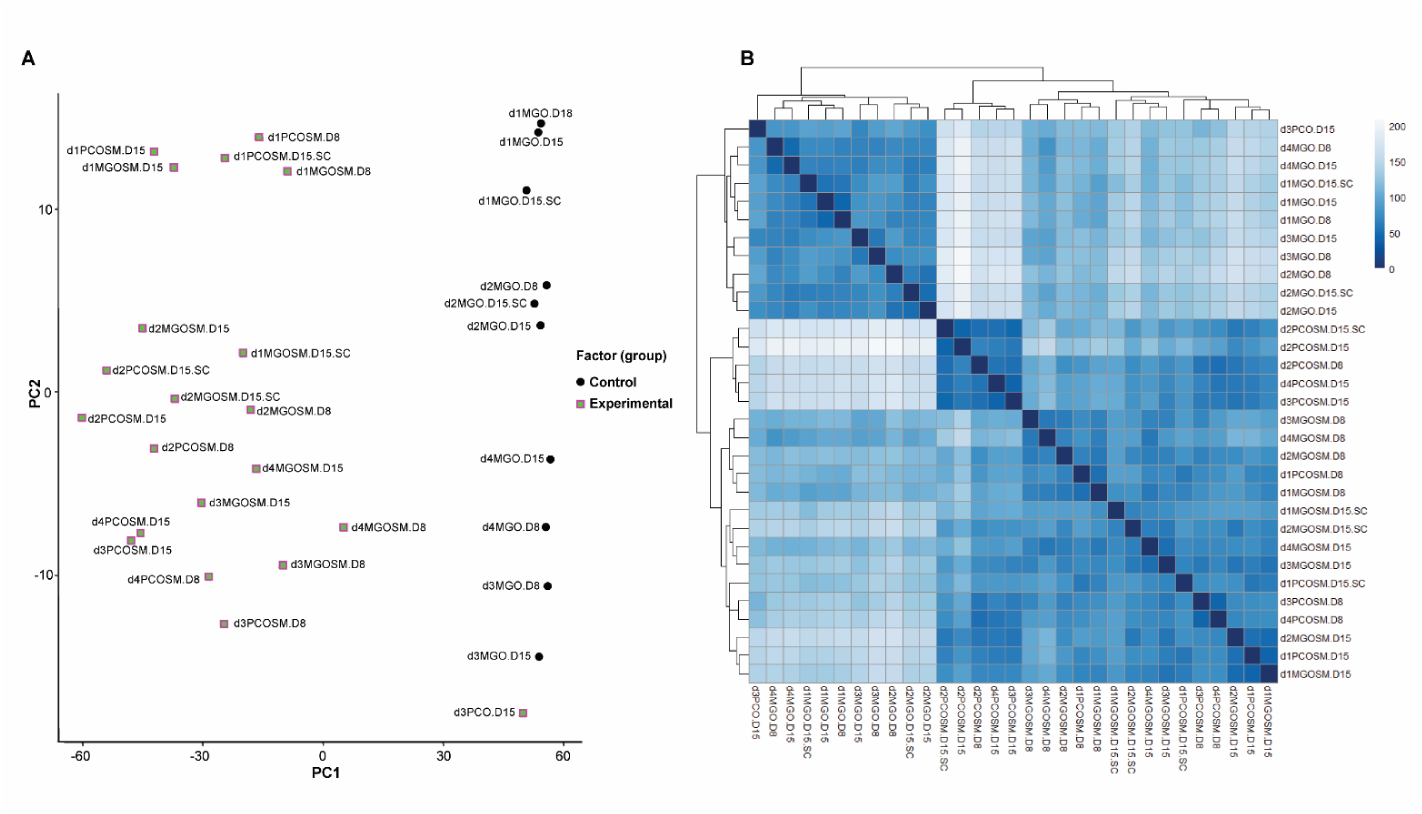
Transcriptomic analysis of all samples used for bulk RNA-seq. (A) PCA graph shows the contribution of the top two principal components to the variance in mRNA expression, with differences between the organoid-only (O) and co-culture (OSM) conditions the most obvious. The second obvious differences are between Matrigel (MG) and PIC+Collagen (PC) hydrogels while the differences between static culture (SC) and dynamic suspension (DS) culture are even less pronounced than donor variations. (B) Pearson correlation among all the samples in the data set shows overall clear separation by influencing factors by cellular complexity (O vs OSM), hydrogels (MG vs PC), donors (d1, d2, d3, d4) and less clear separation by culture conditions (SC vs DS) and differentiation time (D8 vs D15).

**Table S1.**
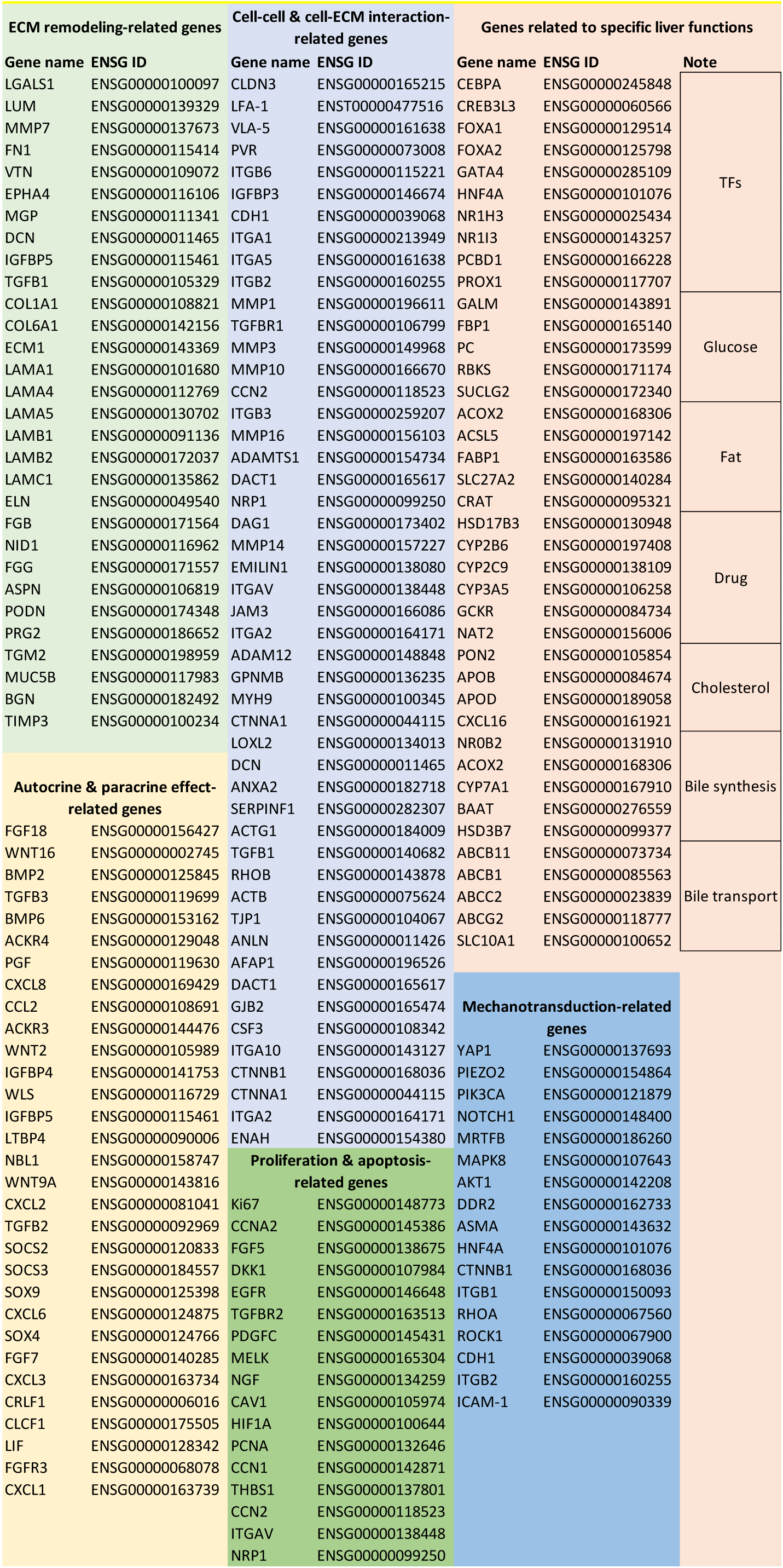
Selected candidate gene lists for heatmap.

**Figure S5.**
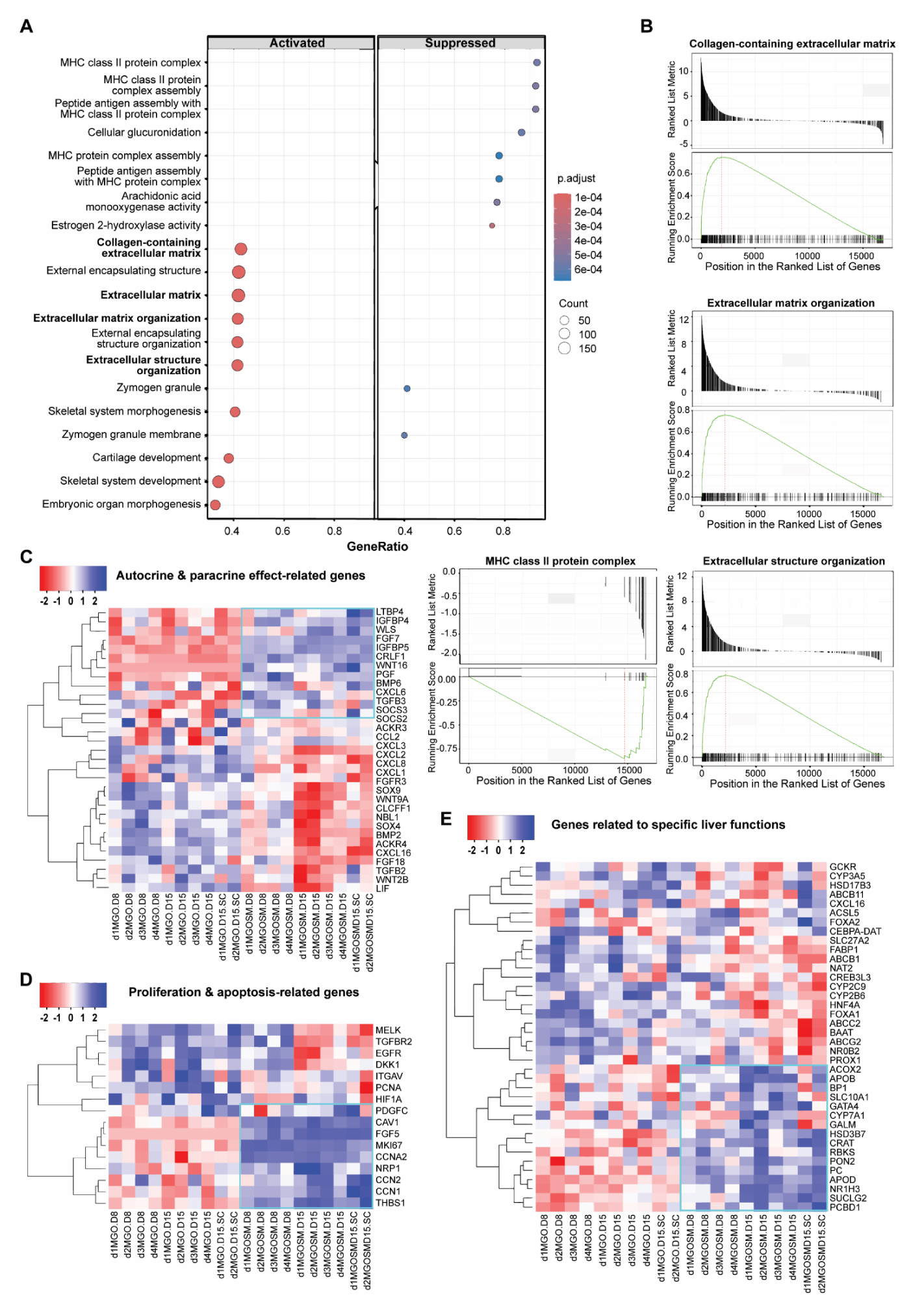
Transcriptomics reveal enhanced ECM remodeling by mesenchymal cells. (A) Gene Ontology (GO) analysis reveals the top involved signaling pathways for the comparison between O and OSM conditions. (B) GSEA plots reveal the potential signaling pathways involved in the functional enhancement of OSM based on the GO gene set. Heatmaps based on the previously reported genes that are associated with autocrine & paracrine effect (C), proliferation & apoptosis (D), or specific liver functions (E).

**Figure S6.**
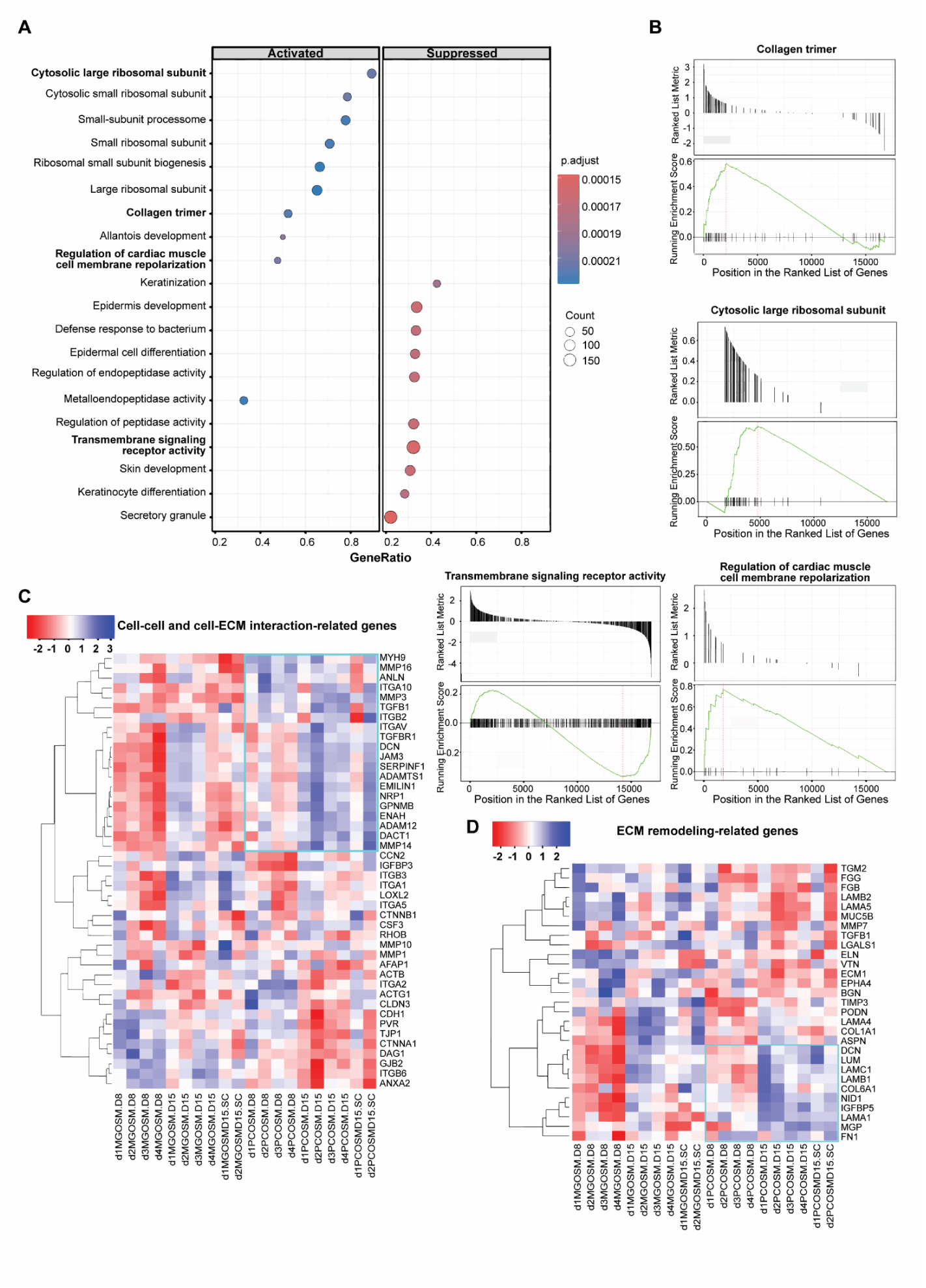
Transcriptomic comparison of BLTs in Matrigel and PC. (A) GO analysis reveals the top involved signaling pathways for all genes comparing MG and PC conditions. (B) GSEA plots reveal the potential signaling pathways involved in the functional enhancement of PC based on the GO gene set. Heatmaps were made out of genes previously reported to be associated with cell-cell & cell-ECM interaction (C) and ECM-remodeling (D).

**Figure S7.**
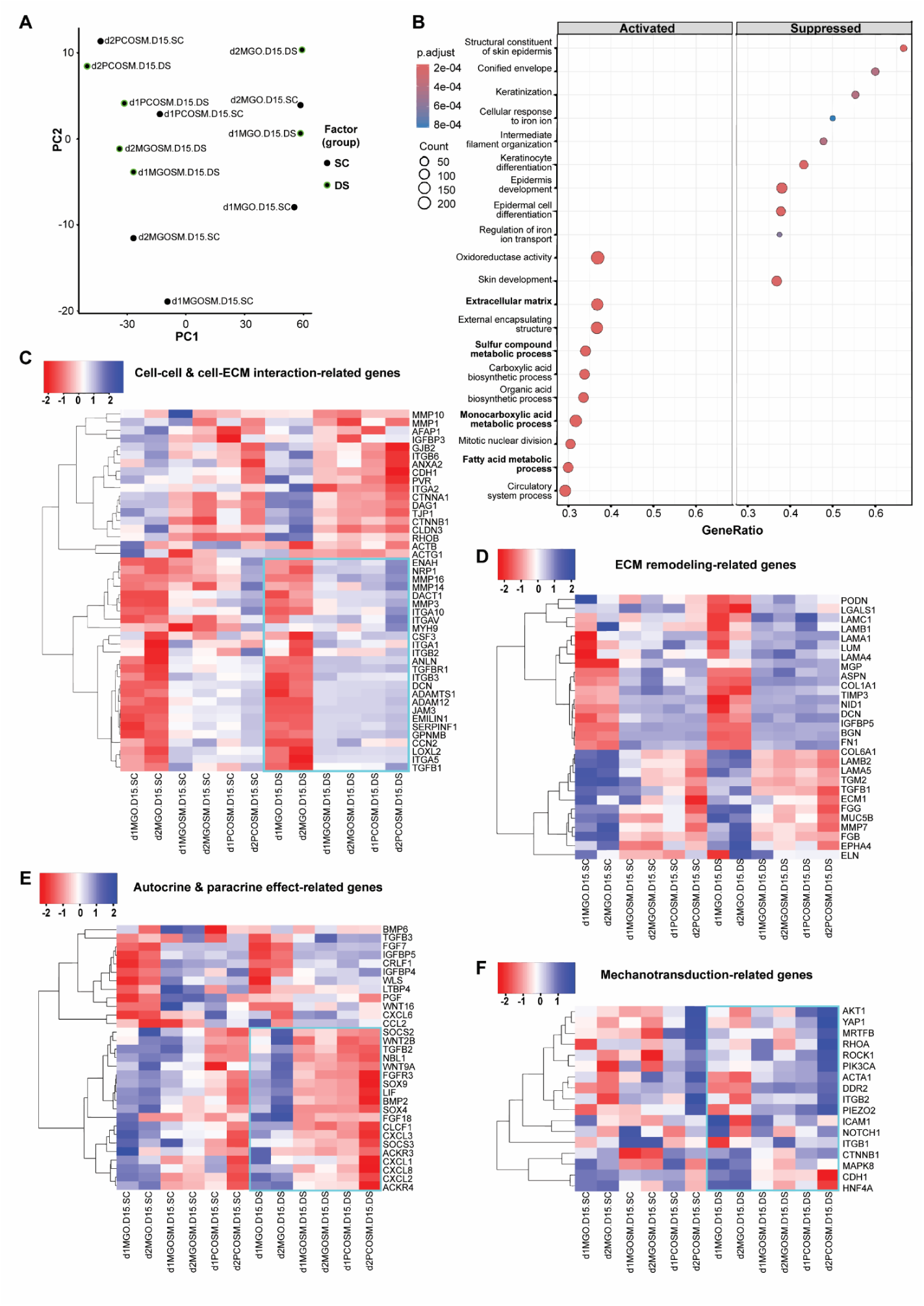
Transcriptomics reveal stronger influences on OSM than on O by dynamic suspension culture. (A) PCA graph shows the contributions of the top two principal components to the variance in mRNA expression between the static culture (SC) and dynamic suspension (DS) conditions with Matrigel (MG) or PIC + Collagen (PC) to form BLTs. The relation between the top two principal components (PC1 and PC2) highlights the separation of samples by culture conditions (SC or DS) and donors (d1 or d2). (B) GO analysis reveals the top involved signaling pathways for all genes comparing SC and DS conditions. Heatmaps were made out of genes previously reported to be associated with cell-cell & cell-ECM interaction (C), ECM-remodeling (D), autocrine & paracrine effect (E), and mechanotransduction (F).

**Figure S8-1.**
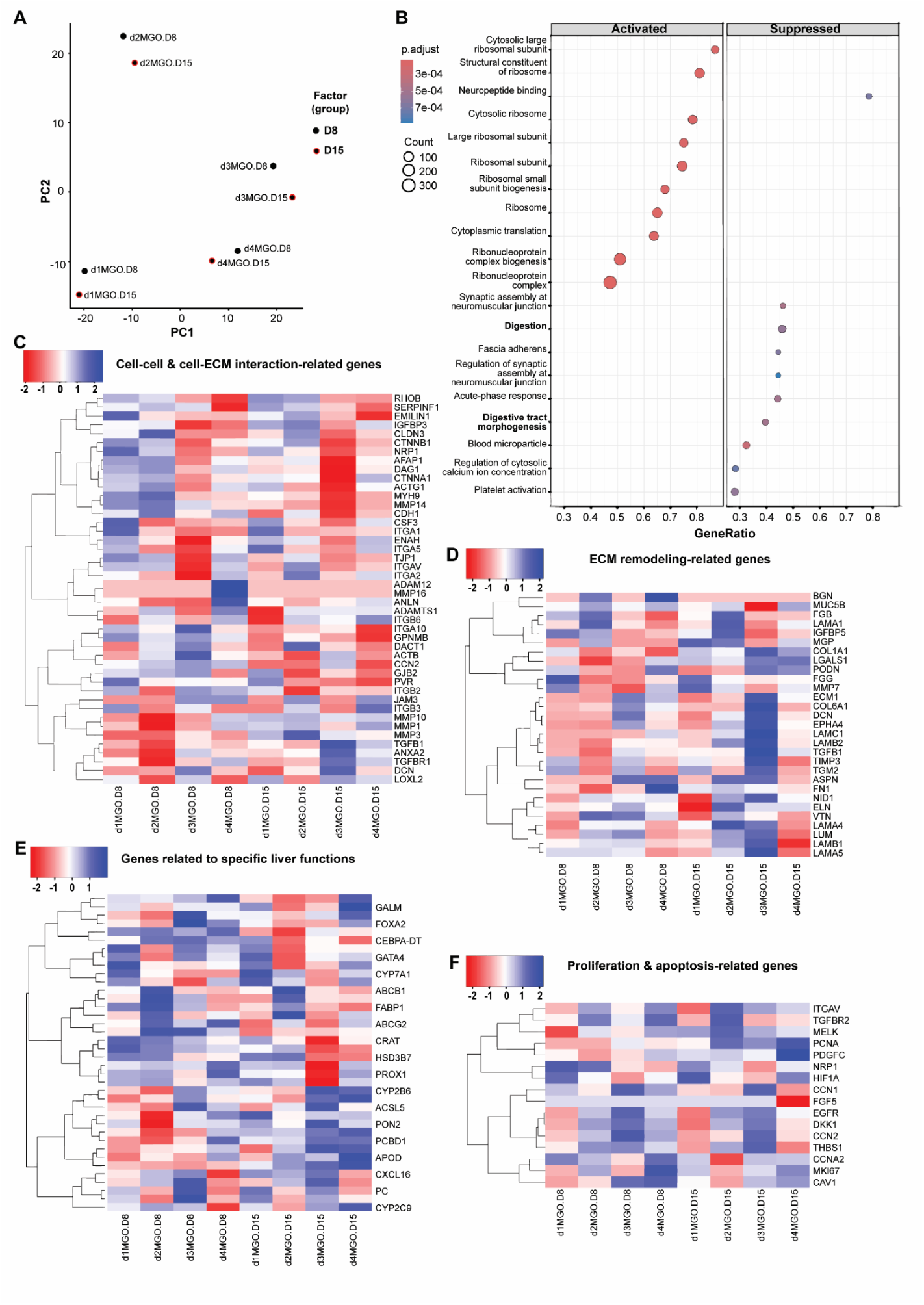
Transcriptomic comparison of organoid-only (O) condition between D8 and D15. (A) PCA graph shows the contributions of the top two principal components to the variance in mRNA expression of BLTs on day 8 (D8) and day 15 (D15) of differentiation. Four donors (d1, d2, d3, d4) were applied. (B) GO analysis reveals the top involved signaling pathways for all genes comparing O samples on D8 and D15. Heatmaps were made as described above with previously reported genes that are associated with cell-cell & cell-ECM interaction (C), ECM-remodeling (D), specific liver functions (E), and proliferation & apoptosis (F).

**Figure S8-2.**
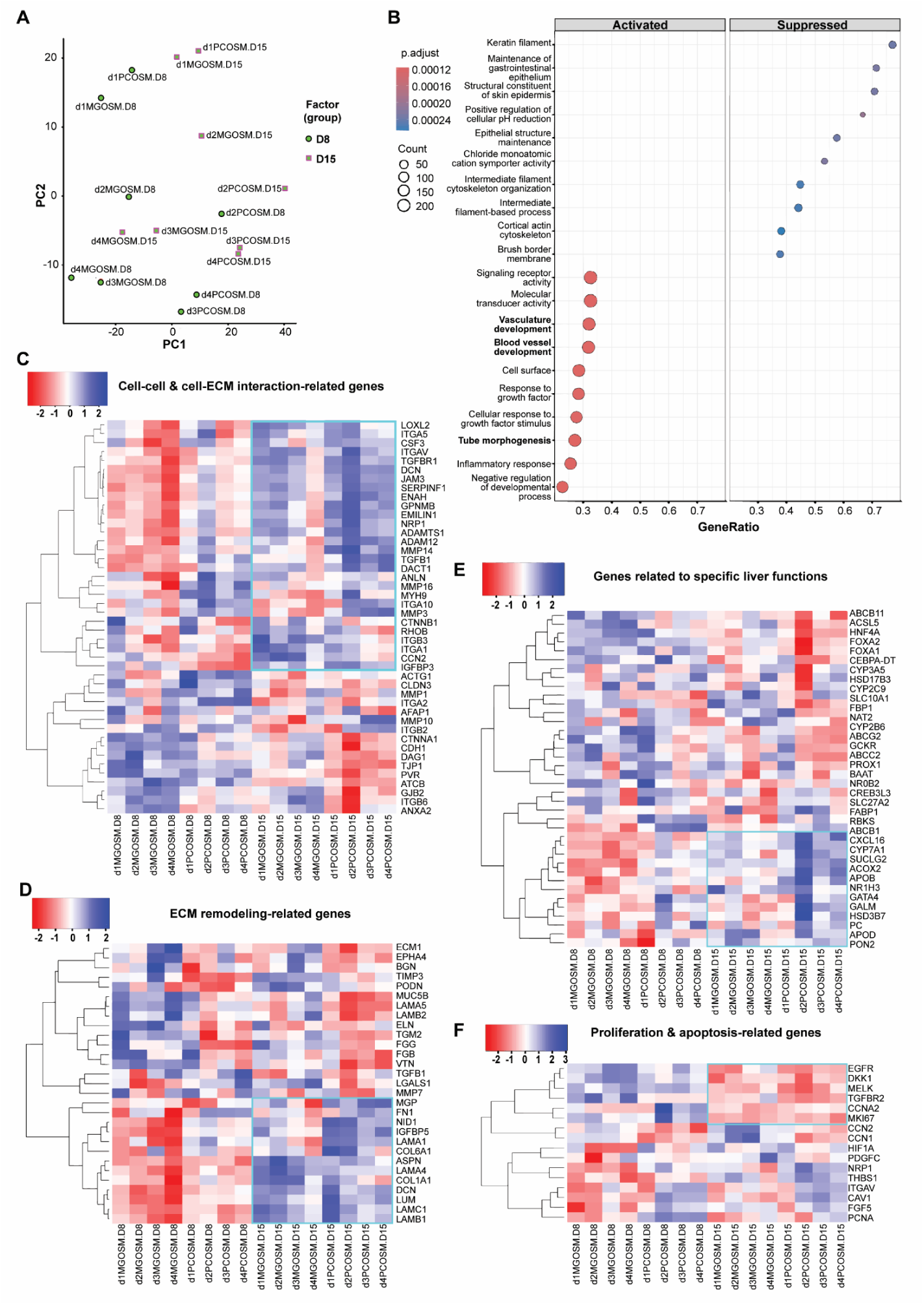
Transcriptomic comparison of the co-culture (OSM) condition between D8 and D15. (A) PCA graph shows the contributions of the top two principal components to the variance in mRNA expression of OSM-derived BLTs on day 8 (D8) and day 15 (D15) of differentiation. Involved factors include two hydrogels, Matrigel (MG) and PIC + Collagen (PC), and four donors (d1, d2, d3, d4). (B) GO analysis reveals the top involved signaling pathways for all genes comparing OSM samples on D8 and D15. Heatmaps were made as described above with previously reported genes associated with cell-cell & cell-ECM interaction (C), ECM-remodeling (D), specific liver functions (E), and proliferation & apoptosis (F).

## Notes

### Competing Interest Statement

The authors have declared no competing interest.

